# The heme biosynthetic enzyme, 5-aminolevulinic acid synthase (ALAS), and GTPases in control of mitochondrial dynamics and ER contact sites, regulate heme mobilization to the nucleus

**DOI:** 10.1101/734780

**Authors:** Osiris Martinez-Guzman, Mathilda M. Willoughby, Arushi Saini, Jonathan V. Dietz, Iryna Bohovych, Amy E. Medlock, Oleh Khalimonchuk, Amit R. Reddi

## Abstract

Heme is an iron-containing cofactor and signaling molecule that is essential for much of aerobic life. All heme-dependent processes in eukaryotes require that heme is trafficked from its site of synthesis in the mitochondria to hemoproteins located throughout the cell. However, the mechanisms governing the mobilization of heme out of the mitochondria, and the spatio-temporal dynamics of these processes, are poorly understood. Herein, using genetically encoded fluorescent heme sensors, we developed a live cell assay to monitor heme distribution dynamics between the mitochondrial inner-membrane, where heme is synthesized, and the mitochondrial matrix, cytosol, and nucleus. We found that heme distribution occurs simultaneously via parallel pathways. In fact, surprisingly, we find that trafficking to the nucleus is ∼25% faster than to the cytosol or mitochondrial matrix. Moreover, we discovered that the heme biosynthetic enzyme, 5-aminolevulinic acid synthase (ALAS), and GTPases in control of the mitochondrial dynamics machinery, Mgm1 and Dnm1, and ER contact sites, Gem1, regulate the flow of heme between the mitochondria and nucleus. Altogether, our results indicate that the nucleus acquires heme faster than the cytosol or mitochondrial matrix, presumably for mitochondrial-nuclear retrograde signaling, and that GTPases that regulate mitochondrial dynamics and ER contact sites are hard-wired to cellular heme distribution systems.

**Summary Statement:** The factors that govern the trafficking of heme, an essential but potentially cytotoxic cofactor and signaling molecule, are poorly understood. Herein, we developed a live-cell assay to monitor heme distribution kinetics and identified the first enzyme in the heme synthesis pathway and GTPases in control of mitochondrial-ER contact sites and dynamics as being critical modulators of heme trafficking.

## Introduction

Heme (iron protoporphyrin IX) is an essential but inherently cytotoxic metallocofactor and signaling molecule (Hanna et al., 2017; Reddi and Hamza, 2016). As a cofactor, heme facilitates diverse processes that span electron transfer, chemical catalysis, and gas synthesis, storage and transport (Hanna et al., 2017; Reddi and Hamza, 2016). As a signaling molecule, heme regulation of an array of proteins, including transcription factors (Ogawa et al., 2001; Pfeifer et al., 1989; Raghuram et al., 2007; Shen et al., 2014), kinases (Chen, 2007; Mense and Zhang, 2006), ion channels (Burton et al., 2016), and micro RNA processing proteins (Barr et al., 2012), collectively control pathways spanning iron homeostasis, oxygen sensing, the oxidative stress response, mitochondrial respiration and biogenesis, mitophagy, apoptosis, circadian rhythms, cell cycle progression, proliferation, and protein translation and degradation (Hanna et al., 2017; Reddi and Hamza, 2016). Although essential for life, heme and its biosynthetic precursors may also act as toxins, necessitating that cells carefully handle the synthesis and trafficking of this compound. The hydrophobicity and redox activity of heme causes it to disrupt membrane structure, become mis-associated with certain proteins, and deleteriously oxidize various biomolecules (Kumar and Bandyopadhyay, 2005; Sassa, 2004). Porphyrin and porphyrinogen, heme biosynthetic intermediates, are photosensitizers that can catalyze the formation of reactive oxygen species like singlet oxygen (Sachar et al., 2016). Indeed, a number of diseases are associated with defects in heme management, including certain cancers (Shen et al., 2014), cardiovascular disease (Wu et al., 2011), aging and age-related neurodegenerative diseases (Atamna and Frey, 2004; Atamna et al., 2002; Schipper et al., 2009), porphyrias (Puy et al., 2010), and anemias (Yang et al., 2016). Despite the tremendous importance of heme in physiology, the cellular and molecular mechanisms that govern the safe assimilation of heme into metabolism remain poorly understood.

In eukaryotes, heme is synthesized via a highly conserved eight-step process. The first and the last three reactions in metazoans (first and last two in the yeast *Saccharomyces cerevisiae*) take place in the mitochondria and the remaining reactions occur in the cytosol (Piel et al., 2019). The first committed step of heme synthesis is the condensation of glycine with succinyl coenzyme A to form 5-aminolevulinic acid (ALA), which is catalyzed by ALA synthase (ALAS). ALA is then exported from the mitochondria into the cytosol where it is converted to coproporphyrinogen III in four steps by the enzymes porphobilinogen synthase (PBGS), hydroxymethylbilane synthase (HMBS), uroporphyrinogen III synthase (UROS), and uroporphyrinogen III decarboxylase (UROD). Coproporphyrinogen III is transported back into the mitochondria where it is converted to protoporphyrin IX (PPIX) in two steps by the enzymes coproporphyrinogen oxidase (CPOX) and protoporphyrinogen oxidase (PPOX). In the final step of heme synthesis, ferrochelatase (FECH) catalyzes the insertion of ferrous iron into PPIX to make heme.

Interestingly, it was recently proposed that the mitochondrial heme biosynthetic enzymes, ALAS, PPOX, and FECH, form a super complex, or metabolon, that includes additional factors like iron and porphyrinogen transporters, and putative heme chaperones (Medlock et al., 2015; Piel et al., 2016). The biochemical rationale for the assembly of a heme metabolon is that it would facilitate the channeling of potentially toxic reactive substrates, including porphyrinogen, porphyrin, and iron, for efficient production and trafficking of heme, thereby mitigating their ability to diffuse freely throughout the cell in an unproductive and deleterious manner. However, given this rationale, it is unclear why ALAS would interact with FECH in the heme metabolon if the product of the ALAS-catalyzed reaction, ALA, were not the immediate substrate for super-complex factors PPOX or FECH. Rather, ALAS may play a regulatory role in heme synthesis or trafficking from FECH beyond simply catalyzing the production of ALA.

Once heme is synthesized by FECH on the matrix side of the mitochondrial inner membrane (IM), it must be mobilized to heme proteins present in virtually every subcellular compartment (Hanna et al., 2017; Reddi and Hamza, 2016). However, the specific factors that govern the transport and trafficking of heme to target hemoproteins are not well-understood (Hanna et al., 2017; Reddi and Hamza, 2016). The current paradigm for subcellular heme trafficking is a *sequential* one in which mitochondrial heme transporters export heme into the cytosol, where it can then be distributed to other locales. The transporter-mediated sequential paradigm for heme distribution was established with the discovery of feline leukemia virus subgroup C cellular receptor 1b (FLVCR1b), the only putative mitochondrial heme transporter identified to date (Chiabrando et al., 2012). FLVCR1b was proposed to transport heme from the mitochondria into the cytosol on the basis that it was required for the hemoglobinization of developing erythrocytes and its ablation results in increased mitochondrial heme content (Chiabrando et al., 2012).

An alternative, but far less studied potential mechanism for heme distribution is through mitochondrial membrane contact sites (Hanna et al., 2017; Piel et al., 2019; Reddi and Hamza, 2016). Endoplasmic reticulum (ER)-mitochondrial encounter structures (ERMES), one type of mitochondrial-ER contact site (Elbaz-Alon et al., 2014), are highly dynamic tethers that physically link the mitochondrial and ER networks and have been proposed to play a role in the transfer of lipids and regulate iron homeostasis (Lackner, 2019; Murley and Nunnari, 2016a; Xue et al., 2017). The frequency of ERMES is in part dependent on mitochondrial division (Elbaz-Alon et al., 2014). ER tubules facilitate mitochondrial fission by wrapping around and constricting mitochondria in an ERMES-dependent fashion, thereby ensuring that the GTPase Dnm1 can oligomerize around a compressed mitochondrial outer-membrane to catalyze the scission of mitochondria (Friedman et al., 2011). Following fission, the GTPase Gem1 disengages ERMES to dissolve the ER-mitochondrial contact site (Murley et al., 2013). As a consequence, ERMES assembly may be dependent on mitochondrial fission and fusion dynamics (Elbaz-Alon et al., 2014). Indeed, mutants with defects in fission have increased ERMES foci, presumably because Dnm1-dependent fission is required for Gem1 to disengage ERMES (Elbaz-Alon et al., 2014). On the other hand, mutants with defects in fusion have decreased ERMES foci (Elbaz-Alon et al., 2014), presumably due to the relative increase in mitochondrial fission over fusion.

Herein, using genetically encoded ratiometric fluorescent <u>h</u>eme <u>s</u>ensors (HS1) targeted to the mitochondrial matrix, cytosol, or nucleus (Hanna et al., 2016; Hanna et al., 2018; Sweeny et al., 2018), we developed a live-cell assay in yeast to monitor heme distribution kinetics to probe the role of ALAS and ER-mitochondrial contact sites and dynamics on subcellular heme trafficking. Surprisingly, we find that heme trafficking rates from the matrix side of the IM to the mitochondrial matrix and cytosol are similar, while trafficking to the nucleus is ∼25% faster. We propose that the increased rate of nuclear heme trafficking is required for heme-based mitochondrial-nuclear retrograde signaling. These data also indicate that heme is distributed from the mitochondrial IM to other locales *simultaneously* via multiple parallel pathways rather than *sequentially.* Moreover, we discovered that the heme biosynthetic enzyme, ALAS, negatively regulates mitochondrial-nuclear heme trafficking, highlighting the close coordination of heme synthesis and trafficking. In addition, we identified GTPases that directly (Gem1) and indirectly (Dnm1 and Mgm1) regulate ERMES as being modulators of nuclear heme transport. Based on our results, we propose a model in which heme is trafficked via ER-mitochondrial membrane contact sites to other organelles such as the nucleus. In total, the development of a live cell assay for probing heme trafficking dynamics coupled with molecular genetics approaches have revealed new insights into subcellular heme distribution.

## Results

### Inter-compartmental heme transport kinetics

In order to probe inter-compartmental heme trafficking in yeast, we developed an *in vivo* pulse-chase assay in which, upon the initiation of heme synthesis, heme mobilization into the mitochondrial matrix, cytosol and nucleus is monitored using genetically encoded ratiometric fluorescent <u>h</u>eme <u>s</u>ensors (HS1) (Hanna et al., 2016). HS1 is a tri-domain fusion protein consisting of a heme-binding moiety, the His/Met coordinating 4-alpha-helical bundle hemoprotein cytochrome *b*_562_ (Cyt *b*_562_), fused to a pair of fluorescent proteins, eGFP and mKATE2, that exhibit heme-sensitive and –insensitive fluorescence, respectively (Hanna et al., 2016) (**Fig. 1a**). Heme binding to the Cyt *b*_562_ domain results in the quenching of eGFP fluorescence via resonance energy transfer but has little effect on mKATE2 fluorescence. Thus, the ratio of eGFP fluorescence (ex: 488 nm, em: 510 nm) to mKATE2 fluorescence (ex: 588 nm, em: 620 nm) reports cellular heme independently of sensor concentration, with the eGFP/mKATE2 ratio inversely correlating with heme binding to the sensor(Hanna et al., 2016). Throughout this study, unless otherwise noted, the sensor is expressed on a centromeric plasmid and the *GPD* promoter (p*GPD*) is used to drive HS1 expression.

**Figure 1.**
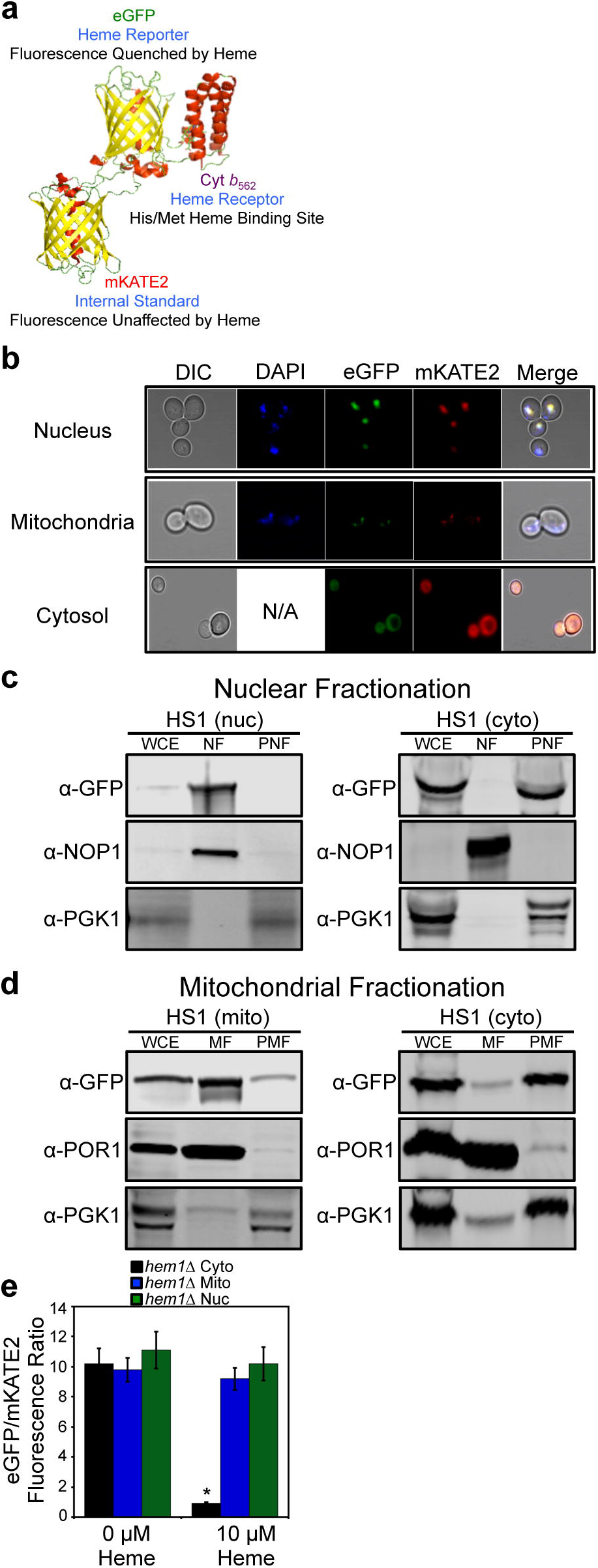
Molecular model and subcellular targeting of heme sensor, HS1. (**a**) Model of HS1 is derived from the X-ray structures of mKATE (PDB: 3BXB) and CG6 (PDB: 3U8P). (**b)** The sub-cellular localization of HS1 was assessed by laser scanning confocal microscopy. Cells expressing nuclear or mitochondrial targeted HS1 were stained with DAPI to label nuclei or mitochondria and live cells were imaged as described in **Materials and Methods** at a magnification of 63x. “Merge” is the merged images of fluorescence from the DAPI, EGFP, and mKATE2 fluorescence channels. (**c** and **d**) Confirmation of mitochondrial (mito), nuclear (nuc), and cytosolic (cyto) HS1 localization as assessed by cell fractionation and immunoblotting. Nuclei (**c**) and mitochondria (**d**) were isolated as described in the **Materials and Methods**, and expression of HS1 was probed using α-GFP antibodies. The indicated fractions were confirmed by probing the expression of PGK1, NOP1, and POR1, which are cytosolic, nuclear, and mitochondrial marker proteins, respectively. 5% of the whole cell extract (WCE), derived from the spheroplast fraction, nuclear fraction (NF), mitochondrial fraction (MF), post-mitochondrial fraction (PMF), or post-nuclear fraction (PNF) were electrophoresed on a 14% tris-glycine SDS-PAGE gel. (**e**) Validation that mitochondrial and nuclear-targeted HS1 does not respond to cytosolic heme. *hem1*Δ cells expressing cytosolic, nuclear, or mitochondrial HS1 were permeabilized with digitonin, a plasma membrane specific permeabilizing agent, and incubated with the indicated concentration of heme. Fluorimetry data represent the mean ± SD of three biological replicates and the statistical significance relative to no heme treatment is indicated by asterisks using an ordinary one-way ANOVA with Dunnett’s post-hoc test. * p < 0.00001.

HS1 is targeted to the mitochondria and nucleus by appending N-terminal Cox4 mitochondrial matrix or C-terminal SV40 nuclear localization sequences, respectively, as previously demonstrated (Hanna et al., 2016) and indicated herein by microscopy (**Fig. 1b**) and organelle isolation and immunoblotting (**Fig. 1c** and **1d**). Indeed, the eGFP and mKATE2 fluorescence emission of nuclear and mitochondrial-targeted HS1 overlay with 4′,6-diamidino-2-phenylindole (DAPI) stained nuclei and mitochondria, respectively (**Fig. 1b**), using a labeling procedure that enables the selective visualization of mitochondrial or nuclear DNA by varying DAPI concentration or exposure time (Shadel and Seidel-Rogol, 2007; Williamson and Fennell, 1979). On the other hand, HS1 lacking nuclear or mitochondrial targeting sequences results in a diffuse pattern of expression throughout the cytosol of the cell. Moreover, isolation of nuclei and mitochondria and immunoblotting for HS1 further confirms the subcellular targeting of mitochondrial and nuclear HS1 (**Fig. 1c** and **1d**). HS1 lacking the nuclear or mitochondrial targeting sequence is found primarily in cytosolic fractions, *i.e.* post-nuclear (PNF) and post-mitochondrial (PMF) fractions, and is not present in significant amounts in mitochondrial or nuclear fractions (**Fig. 1c** and **1d**).

It is possible that a small fraction of mitochondrial or nuclear-targeted sensor may be present in the cytosol and confound analysis of sub-compartmental heme. In order to assess if this were the case, we permeabilized heme-deficient cells lacking *HEM1*, which encodes the first enzyme in the heme biosynthetic pathway, ALAS, with digitonin, a mild non-ionic detergent that selectively permeabilizes the plasma membrane but not mitochondrial or nuclear membranes, and treated cells with heme. As indicated in **Fig. 1e**, only cells expressing cytosolic HS1 exhibited a significant heme-dependent reduction in eGFP/mKATE2 fluorescence ratio, with no significant perturbations to the fluorescence of mitochondrial and nuclear-targeted HS1. These data indicate that mitochondrial and nuclear-targeted HS1 is unresponsive to cytosolic heme. Thus, altogether, our data indicate that cytosolic, nuclear, and mitochondrial heme can be robustly monitored *in vivo* using HS1.

Inter-compartmental heme trafficking rates are monitored by: **a.** inhibiting heme synthesis with 500 μM succinylacetone (SA), an inhibitor of PBGS, for ∼16 hours in sensor expressing cells; **b.** removing the block in heme synthesis by re-suspending cells into media lacking SA; and **c.** monitoring the time-dependent change in the eGFP/mKATE2 ratio (R) of HS1 localized to different locations upon the re-initiation of heme synthesis (**Fig. 2a-d**). As described in the **Materials and Methods** section and **Equation 1**, the fractional saturation of HS1 (% Bound) is calculated by considering R relative to the HS1 fluorescence ratio when the sensor is 0% bound (*R*_min_) and 100% bound (*R*_max_) (**Fig. 2e**), which are parameters derived from parallel cultures of cells continually maintained in media with and without SA (Hanna et al., 2016; Hanna et al., 2018) (**Fig. 2a-c**).

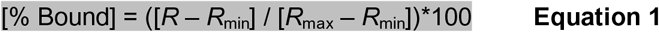

**Figure 2.**
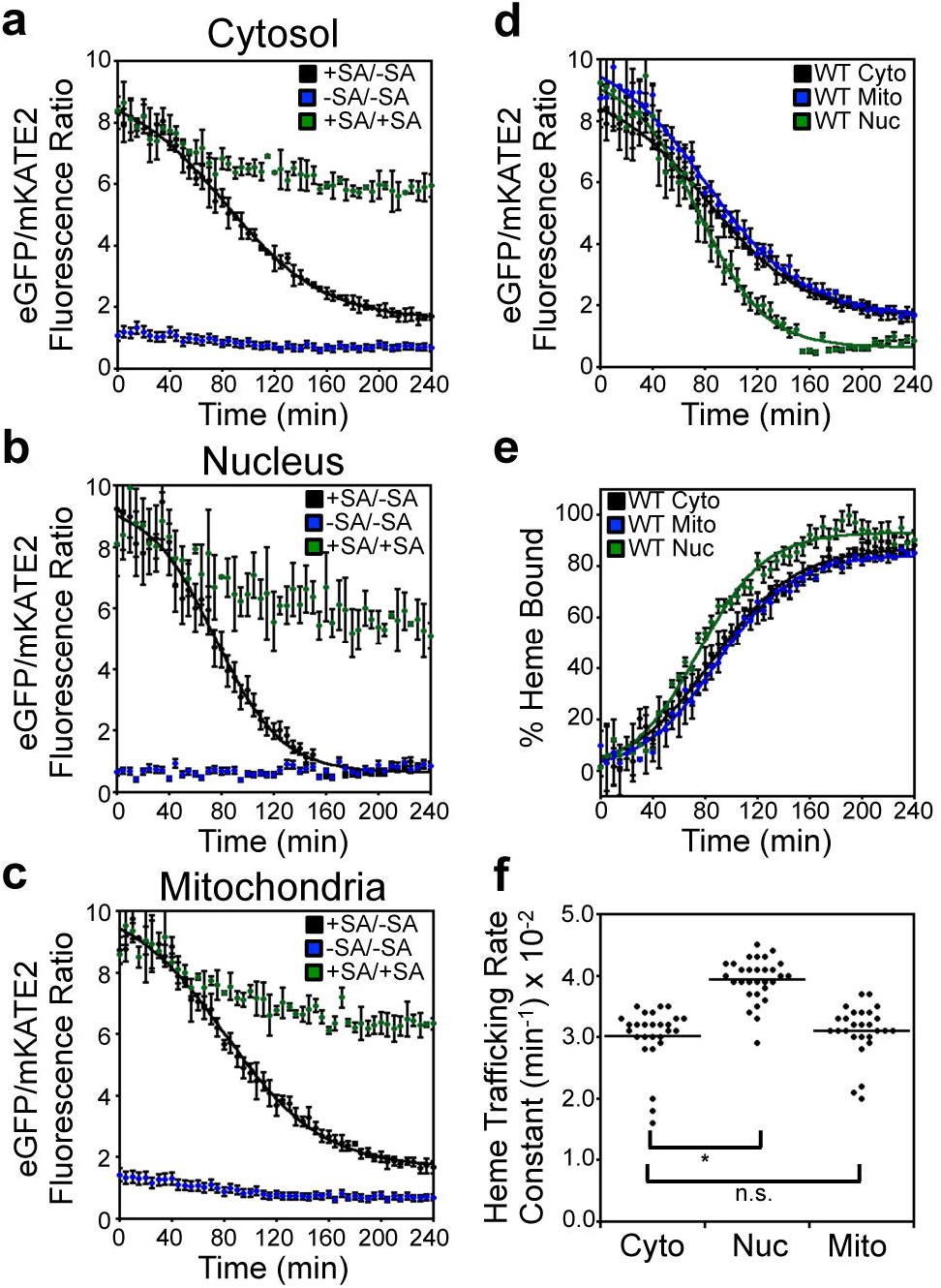
Inter-compartmental heme trafficking dynamics as measured by heme sensor HS1. WT cells expressing HS1 in the cytosol (**a**), nucleus (**b**), or mitochondria (**c**) were depleted of heme using 500 μM succinylacetone (SA), the heme biosynthetic inhibitor and, upon the re-initiation of heme synthesis, the rates of heme trafficking to the indicated subcellular locations were monitored by measuring the time-dependent change in sensor eGFP/mKATE2 fluorescence ratio (**d**) or the fractional saturation of HS1 (**e**), which is determined using **equation 1** and the “raw” fluorescence ratio values in **a**-**c**. The data in **d** and **e** are fit to **equation 2**. (**f**) The rate constants for cytosolic, nuclear, and mitochondrial heme trafficking, which were extracted from fits to the data represented in panel “**e**”, from triplicate cultures in nine independent experimental trials are shown. The statistical significance relative to the rate constants for cytosolic heme trafficking is indicated by an asterisk using an ordinary one-way ANOVA with Dunnett’s post-hoc test. * p < 0.0001. The heme trafficking kinetics data shown in all figure panels represent the mean ± SD of independent triplicate cultures.

In order to confirm the SA-mediated block in heme synthesis, we determined that SA-conditioned wild type (WT) cells exhibit intracellular heme concentrations (Michener et al., 2012) (**Fig. S1a**) and HS1 eGFP/mKATE2 ratios (Hanna et al., 2016) (**Fig. S1b**) similar to *hem1*Δ cells, which cannot make heme due to the deletion of the first enzyme in the heme biosynthetic pathway, ALAS (Hanna et al., 2016). SA pre-conditioned cells shifted to media lacking SA exhibit a time-dependent decrease in HS1 eGFP/mKATE2 ratio (**Fig. 2a-c**) in the cytosol, nucleus, and mitochondria, which is characteristic of increased heme loading of the sensor due to newly synthesized heme (**Fig. 2e**). By contrast, the HS1 eGFP/mKATE2 ratios in cells continuously maintained with or without SA remain characteristically high or low, respectively (**Fig. 2a-c**), albeit there are some modest time dependent changes for currently unknown reasons.

The change in sensor heme saturation (% Bound) following the re-initiation of heme synthesis reveals three distinct phases: a lag phase, an exponential phase, and a stationary phase (**Fig. 2e**). The lag phase can be interpreted as the time required to alleviate the SA-mediated block in heme synthesis and re-initiate heme production. The exponential phase represents the rate of heme binding to HS1, which is governed by the relative rates of heme synthesis and trafficking to the different subcellular locations. However, when comparing HS1 heme saturation kinetics between different compartments within a given strain, the data strictly represent the inter-compartmental heme trafficking rates since the contribution from heme synthesis is a constant. Indeed, expression of cytosolic, nuclear, or mitochondrial HS1 does not perturb heme synthesis (**Fig. S2a**). Since the final step of heme synthesis occurs with the insertion of ferrous iron into protoporphyrin IX, a reaction catalyzed by FECH at the matrix side of the mitochondrial IM, the heme trafficking rates reflect heme mobilization from the mitochondrial IM to the matrix, cytosol, or nucleus. Given the high affinity of HS1 for both ferric and ferrous heme, *K*_D_^III^ = 10 nM and *K*_D_^II^ < 1 nM at pH 7.0(Hanna et al., 2016), and the negligible differences in FRET between eGFP and ferric and ferrous heme(Hanna et al., 2016), we cannot resolve the two oxidation states of heme. The stationary phase represents the maximum limiting heme saturation of the sensor 4 hours after alleviating the SA-mediated block in heme synthesis and typically spans ∼70-100%.

The kinetics of heme binding to HS1 (**Fig. 2e**) can be fit to the logistic function described in **Equation 2**, where [A] is the maximal value of HS1 fractional saturation (amplitude), *k* is the first order rate constant (min^-1^), *t* is the time (min), and *t*_1/2_ is the midpoint of the transition. The lag time can be defined as *t*_1/2_ – 2/*k* (Arosio et al., 2015).

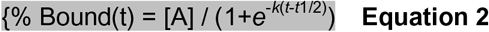

Quite surprisingly, the kinetics of HS1 heme saturation indicate that heme trafficking to locations as disparate as the mitochondrial matrix, cytosol, and nucleus are similar, but with heme transport to the nucleus exhibiting a ∼25% increase in rate constant relative to heme transport to the cytosol and mitochondrial matrix (**Fig. 2e** and **2f**). This observation is highly reproducible and the rate constants for heme trafficking to the cytosol (*k*^CYTO^), nucleus (*k*^NUC^), and mitochondrial matrix (*k*^MITO^) were found to be 0.030±.005 min^-1^, 0.039±.003 min^-1^, and 0.031±.004 min^-1^, which are the average ± standard deviation from triplicate biological samples over nine independent experimental trials (**Fig. 2f**). Importantly, in every trial conducted, even if the absolute rates were different, the relative differences always followed the same trend, *i.e. k*^NUC^ > *k*^CYTO^ ∼ *k*^MITO^. In addition, fitting of the “raw” unprocessed sensor ratios, *R*, reveal a similar trend in the relative rates of heme binding to HS1, with *k*^NUC^ (0.039±.003 min^-1^) > *k*^CYTO^ (0.026±.001 min^-1^) = *k*^MITO^ (0.027±.002 min^-1^) (**Fig. 2d**), albeit the absolute rate constants are different when fitting processed “% Bound” traces. This is due to the fact that, for reasons not completely understood, there are fluctuations in *R*_min_ and *R*_max_ over the course of a heme trafficking dynamics experiment (**Fig. 2a-c**). Although the qualitative trends are similar between analyzing changes in “% Bound” and “*R*”, “% Bound” is the preferred metric since it accounts for non-heme dependent changes in sensor fluorescence ratio, *i.e.* fluctuations in *R*_min_ or *R*_max_ over the course of an experiment.

The remarkable similarity in the observed rates of heme trafficking to different locales might suggest that heme binding to the sensor is rate limiting. However, measurements of the rate of heme binding to HS1 in cell lysates of WT cells depleted of heme indicate that heme binding to the cytosolic, mitochondrial, and nuclear sensors occurs in less than 2 minutes (**Fig. S3**), which is much faster than the ∼90-120 minutes it takes 50% of: (i) total cellular heme to be synthesized (**Fig. S2a**) or (ii) HS1 to become heme saturated in the SA pulse-chase assays (**Fig. 2**).

In order to rule out any potential artifacts associated with heme sensor expression itself perturbing the observed heme trafficking rates, we expressed cytosolic HS1 under the control of weak (p*ADH1*), medium (p*TEF1*), or strong (p*GPD*) promoters and found that the observed heme trafficking rates were not affected despite a nearly 10-fold span in sensor expression (**Fig. S2b**). These results are consistent with our previous findings that sensor expression does not perturb heme homeostasis or otherwise affect cell viability(Hanna et al., 2016). Analogous experiments titrating expression with mitochondrial and nuclear sensors could not be completed due to low sensor expression and correspondingly low sensor signal to noise ratios associated with p*TEF1*-or p*ADH1*-driven expression. Altogether, our data indicate that the SA-pulse chase assays can be used to measure the rates of heme trafficking to multiple compartments and that the rates of mitochondrial-nuclear heme trafficking are 25% faster relative to heme trafficking to the cytosol and mitochondria.

In order to rule out that the observed rates of inter-compartmental heme trafficking are due to artifacts associated with SA treatment, we re-capitulated the *in vivo* heme transport assay in *hem1*Δ cells pulsed with a bolus of ALA to initiate heme synthesis. As shown in **Fig. 3a**, heme trafficking kinetics to the matrix, cytosol, and nucleus in *hem1*Δ cells are qualitatively similar to the results obtained from the SA pulse-chase assay using WT cells (**Fig. 1e** and **1f**), with transport to the nucleus (*k*^NUC^ = 0.059±.005 min^-1^) being faster than to the cytosol (*k*^CYTO^ = 0.030±.003 min^-1^) and mitochondrial matrix (*k*^MITO^ = 0.028±.004 min^-1^) (**Table S1**). Notably, the rate enhancement for heme trafficking to the nucleus is greater in magnitude in *hem1*Δ cells pulsed with ALA (*k*^NUC^ / *k*^CYTO^ = 2.0 ± 0.1; **Fig. 3a**) than in WT cells that have the SA-mediated block in heme synthesis alleviated (*k*^NUC^ / *k*^CYTO^ = 1.3 ± 0.2); **Fig. 2e** and **2f**), one-way ANOVA, p < .01, n = 3. This observation suggests that either the use of SA suppresses nuclear heme trafficking or that the first enzyme in the heme biosynthetic pathway, ALAS, is a negative regulator of mitochondrial-nuclear heme trafficking.

**Figure 3.**
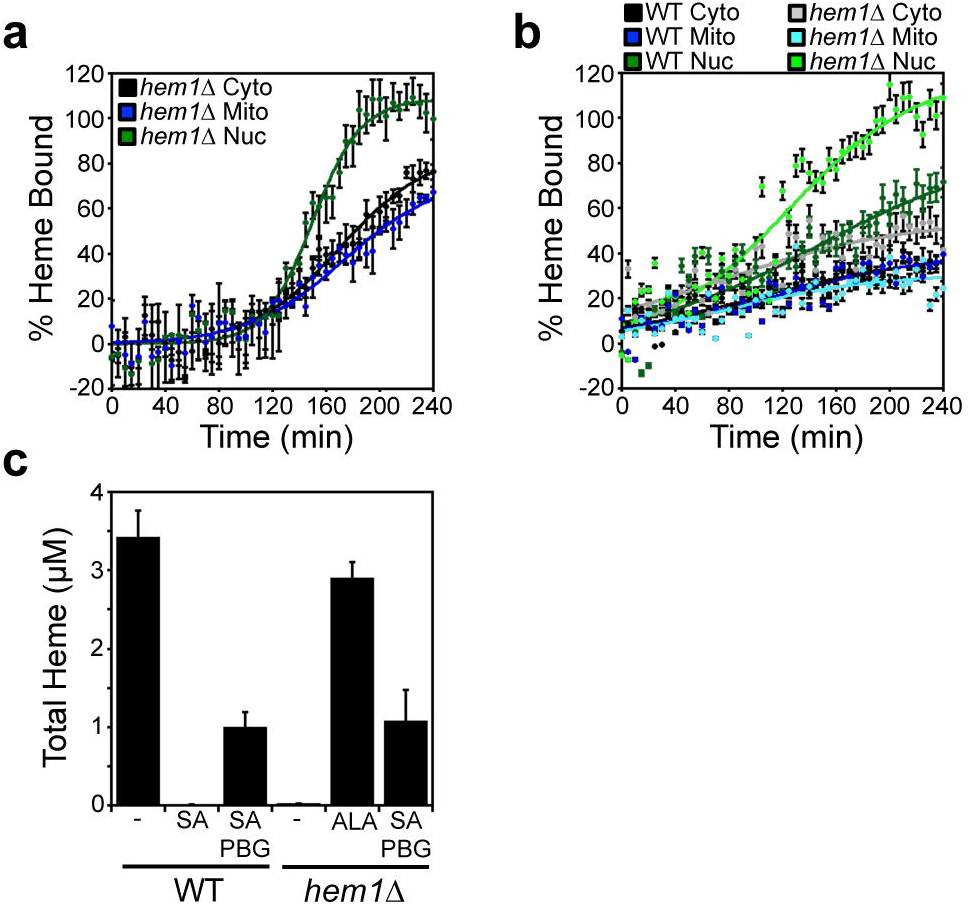
ALA synthase (Hem1) negatively regulates mitochondrial-nuclear heme trafficking. (**a**) *hem1*Δ cells expressing HS1 in the cytosol (black), nucleus (green), or mitochondria (blue) were pulsed with a bolus of ALA to initiate heme synthesis and the rates of heme trafficking to the indicated subcellular locations were monitored by measuring the fractional saturation of HS1 over time as in **Fig. 1e**. Fits to the data using **equation 2**, reveal *k*^CYTO^ = 0.030±.003 min^-1^, *k*^NUC^ = 0.059±.005 min^-1^ and *k*^MITO^ = 0.028±.004 min^-1^, which are averages and standard deviations from three independent cultures. (**b**) WT and *hem1*Δ cells expressing sensor were pre-cultured with succinylacetone (SA) to deplete heme and pulsed with a bolus of porphobilinogen (PBG) to re-initiate heme synthesis. The rates of heme trafficking to the indicated subcellular locations were monitored by measuring the fractional saturation of HS1 over time as in **Fig. 1e**. Fits to the data using **equation 2**, reveal WT trafficking rate constants to be *k*^CYTO^ = 0.012±.002 min^-1^ (black), *k*^NUC^ = 0.018±.002 min^-1^ (dark green), and *k*^MITO^ = 0.013±.002 min^-1^ (dark blue), and *hem1*Δ trafficking rate constants to be *k*^CYTO^ = 0.014±.003 min^-1^ (grey), *k*^NUC^ = 0.028±.003 min^-1^ (light green), and *k*^MITO^ = 0.012±.002 min^-1^ (light blue). The indicated rate constants and data are the mean and standard deviations from independent triplicate cultures. (**c**) Analysis of total heme in WT and *hem1*Δ strains treated with succinylacetone (SA), aminolevulinic acid (ALA), or porphobilinogen (PBG). The data represent the mean ± SD of duplicate cultures.

### ALA synthase (ALAS) regulates mitochondrial-nuclear heme trafficking

In order to determine if ALAS plays a role in regulating heme distribution kinetics, we monitored inter-compartmental heme trafficking kinetics directly between WT and *hem1*Δ cells. Towards this end, we monitored heme distribution dynamics in WT and *hem1*Δ cells that had SA-inactivated PBGS pulsed with 500 μM porphobilinogen (PBG), the product of the reaction catalyzed by PBGS, to re-initiate heme biosynthesis. As with the SA-pulse chase experiment in WT cells (**Fig. 2e** and **2f**), the rates of heme trafficking to the nucleus are greater than trafficking to the cytosol and mitochondrial matrix in SA-treated WT cells pulsed with PBG (**Fig. 3b**); *k*^NUC^ / *k*^CYTO^ = 1.5 ± 0.2 vs. *k*^MITO^ / *k*^CYTO^ = 1.1 ± 0.2; one-way ANOVA, p = 0.05, n = 3. Strikingly, in SA-treated *hem1*Δ cells pulsed with PBG, heme trafficking to the nucleus is significantly augmented relative to WT cells (**Fig. 3b**); *k*^NUC^ / *k*^CYTO^ = 2.0 ± 0.1 in *hem1*Δ cells vs. *k*^NUC^ / *k*^CYTO^ = 1.5 ± 0.2 in WT cells, one-way ANOVA, p = 0.05, n = 3. The relatively low final fractional saturation of the sensor using PBG can be attributed to the fact that heme-deprived cells are less efficient at using PBG to make heme (**Fig. 3c**). WT and *hem1*Δ cells treated with SA and PBG make roughly ∼33% of the heme that WT or *hem1*Δ cells treated with ALA make. Thus, it is all the more impressive that in *hem1*Δ cells pulsed with PBG, the nuclear sensor is completely saturated with heme, unlike in WT cells. Importantly, *hem1*Δ cells do not have a defect in synthesizing heme when supplied with PBG (**Fig. 3c**). Thus, in total, our results demonstrate that ALAS negatively regulates mitochondrial-nuclear heme trafficking without impacting downstream steps of heme synthesis.

### Gem1, a GTPase that regulate ERMES, negatively modulates mitochondrial-nuclear heme trafficking

Since the final step of heme synthesis occurs on the matrix side of the mitochondrial IM, our expectation was that heme would sequentially populate sensors in the mitochondrial matrix, cytosol, and nucleus. However, the data indicate that once heme is synthesized in the mitochondrial IM, it disperses to multiple compartments almost simultaneously, suggesting the existence of parallel routes for heme mobilization to distinct locales. Moreover, surprisingly, mitochondrial-nuclear heme trafficking occurs ∼25% faster than to other compartments. Since mitochondria form dynamic physical contacts with other organelles like the ER (Elbaz-Alon et al., 2014; Murley and Nunnari, 2016a), we surmised that one potential route for the distribution of mitochondrial heme to other compartments like the nucleus could be through such membrane contact sites. In this scenario, heme may be trafficked to the nucleus via the ER network, bypassing the cytosol, thereby accounting for the increased rate of nuclear heme trafficking. To test this idea, we probed heme distribution rates in deletion mutants with increased or decreased ER-mitochondrial contact sites.

The ER-mitochondrial encounter structure (ERMES), which constitutes one type of physical interface between these organelles, is a complex that consists of four proteins, Mmm1, Mdm10, Mdm12, and Mdm34 (Boldogh et al., 2003; Kornmann et al., 2009; Sogo and Yaffe, 1994; Youngman et al., 2004). Mmm1 and Mdm10 are anchored in the ER and outer mitochondrial membranes, respectively, while the cytosolic protein Mdm12 and the mitochondrial protein Mdm34 help to bridge the interaction between Mmm1 and Mdm10 in the ERMES complex (Boldogh et al., 2003; Kornmann et al., 2009; Sogo and Yaffe, 1994; Youngman et al., 2004). The absence of any single ERMES protein results in dissociation of the entire complex, compromising the formation of ER-mitochondrial contact sites, mitochondrial division and morphology(Boldogh et al., 2003; Burgess et al., 1994; Kornmann et al., 2009; Sogo and Yaffe, 1994; Youngman et al., 2004). ERMES is negatively regulated by Gem1, which is a mitochondrial outer-membrane localized GTPase that disengages ERMES after mitochondrial division(Kornmann et al., 2011; Murley et al., 2013).

As shown in **Figure 4a**, *gem1*Δ cells, which have increased ERMES, exhibit a significantly increased rate of mitochondrial-nuclear heme trafficking, with minimal impact on heme trafficking to the cytosol and mitochondrial matrix; *k*^NUC^ / *k*^CYTO^ = 1.45 ± 0.05 in *gem1*Δ cells vs. *k*^NUC^ / *k*^CYTO^ = 1.1 ± 0.1 in WT cells, one-way ANOVA, p < 0.01, n = 3. These data suggest that more stable ER-mitochondrial contacts increase the flow of heme to the nucleus. However, in ERMES compromised *mdm12*Δ and *mdm10*Δ strains, heme trafficking rates are unaffected (**Fig. 4b** and **4c**).

**Figure 4.**
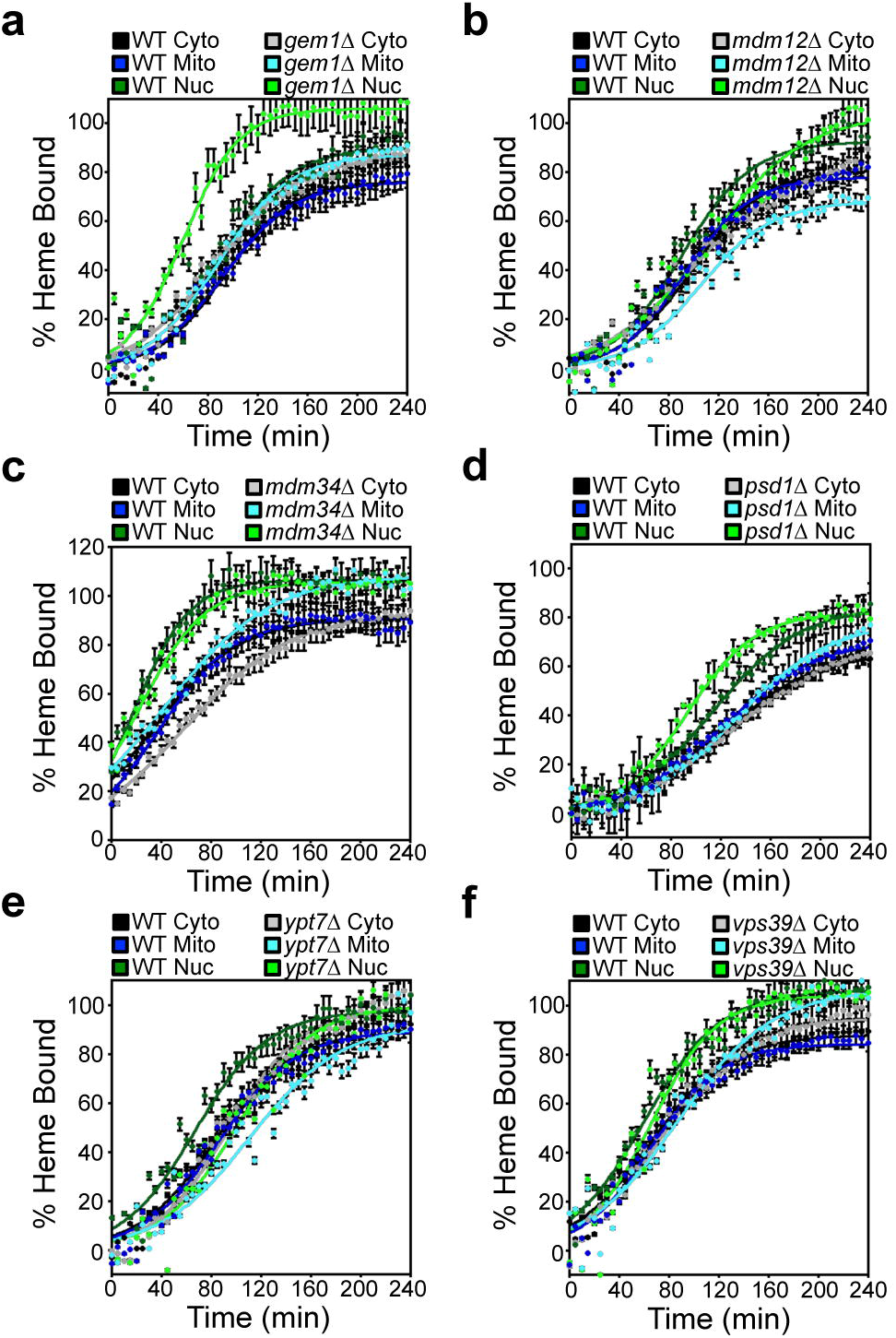
Gem1 is a negative regulator of mitochondrial-nuclear heme trafficking. Inter-compartmental heme trafficking rates were monitored as in **Fig. 1c** using the SA pulse-chase assay for (**a**) *gem1*Δ, (**b**) *mdm12*Δ, (**c**) *mdm34*Δ, (**d**) *psd1*Δ, (**e**) *ypt7*Δ, and (**f**) *vps39*Δ cells. The data represent the mean ± SD of independent triplicate cultures and the kinetic parameters derived from the fits to the data using equation 2 are indicated in **Table S1**.

ERMES has been implicated in the transfer of lipids, *e.g.* phosphatidylserine (PS), between the ER and mitochondria (Kawano et al., 2018; Kornmann et al., 2009; Nguyen et al., 2012). PS, an ER-synthesized lipid, is a precursor for the synthesis of phosphatidylethanolamine (PE) in the mitochondria, which itself is a precursor for phosphatidylcholine (PC) synthesis back in the ER. Both phospholipids are constituents of mitochondrial membranes (Schuler et al., 2016). In order to rule out confounding contributions that may arise due to general defects in PE or PC in ERMES mutants, we tested heme trafficking dynamics in *psd1*Δ cells, which lack PS decarboxylase, the enzyme that catalyzes the formation of PE. As shown in **Fig. 4d**, there are no defects in heme trafficking dynamics in *psd1*Δ cells.

ERMES are highly dynamic and sensitive to the presence of mitochondrial-vacuolar contact sites, termed vCLAMPs for vacuolar and mitochondrial patches. Like ERMES, vCLAMPs are important for lipid and metabolite trafficking between mitochondria and the endomembrane system (Elbaz-Alon et al., 2014). The absence of one causes expansion of the other and ablation of both ERMES and vCLAMPs is lethal (Elbaz-Alon et al., 2014). We therefore sought to determine if a defect in vCLAMPs also increases mitochondrial-nuclear heme trafficking rates as in *gem1*Δ cells, which have an increased number of ERMES foci like vCLAMP mutants. However, neither *vps39*Δ nor *ypt7*Δ cells (**Fig. 4e** and **4f**), which lack a component of the HOPS (‘homotypic fusion and protein sorting’) tethering complex (Vps39) and the Rab GTPase (Ypt7), core components of vCLAMPs (Elbaz-Alon et al., 2014; Honscher et al., 2014; Murley and Nunnari, 2016b), exhibit defects in heme distribution kinetics. We conclude that Gem1, but not the core components of ERMES and vCLAMP machineries, regulates mitochondrial-nuclear heme trafficking.

### Mgm1 and Dnm1 are positive and negative regulators of mitochondrial-nuclear heme trafficking, respectively

The mitochondrial network is highly dynamic and is constantly remodeled by fusion and fission events. The dynamic behavior of the mitochondrial network is thought to be responsible for the proper cellular distribution and trafficking of a number of mitochondrial-derived metabolites, including various lipids (Chan, 2012; Tatsuta et al., 2014). Given that we identified Gem1, an ERMES regulating GTPase that modulates mitochondrial-nuclear heme trafficking, and that ERMES are associated with sites of mitochondrial division and their frequency can be impacted by mitochondrial fission and fusion dynamics (Elbaz-Alon et al., 2014; Friedman et al., 2011), we reasoned that other GTPases that regulate mitochondrial dynamics might regulate mitochondrial-nuclear heme trafficking.

#### Mitochondrial Fusion

Mitochondrial fusion occurs at both the IM and OM in coordinated but physically separable steps. In yeast, a pair of dynamin-like GTPases, Fzo1 and Mgm1, drive OM and IM fusion, respectively (Westermann, 2008). In order to coordinate double membrane fusion, a protein spanning the mitochondrial intermembrane space, Ugo1, physically tethers Fzo1 and Mgm1. Mgm1 exists in equilibrium between long (l-Mgm1) and short (s-Mgm1) isoforms, which are generated upon proteolytic cleavage by the rhomboid protease Pcp1 (DeVay et al., 2009; Zick et al., 2009). It is thought that l-Mgm1, which has an inactive GTPase domain, acts as an anchor in the IM and interacts with and activates the GTPase activity of s-Mgm1 in the intermembrane space to promote IM fusion (DeVay et al., 2009; Zick et al., 2009).

Relative to WT cells, the *mgm1*Δ strain exhibits a marked defect in nuclear heme trafficking, with a > 50% reduction in HS1 heme loading (**Fig. 5a**). On the other hand, heme trafficking to the mitochondrial matrix and cytosol is not significantly impacted in *mgm1*Δ cells. Interestingly, heme availability to HS1 in the nucleus is similar between WT and *mgm1*Δ cells after prolonged growth, ∼16 hours, suggesting that there are compensatory mechanisms that overcome the defects in nuclear heme trafficking in *mgm1*Δ cells (**Fig. S4a**). The rate of heme synthesis is un-perturbed by loss of Mgm1, indicating that the nuclear trafficking defect is not related to general defects in heme synthesis (**Fig. S4b**). The effects of the *mgm1*Δ mutant on nuclear heme trafficking is not due to the loss of mitochondrial DNA, which occurs with high frequency in *mgm1*Δ cells; *rho*^0^ cells that lack mitochondrial DNA do not exhibit an appreciable defect in nuclear heme trafficking (**Fig. 5b**).

**Figure 5.**
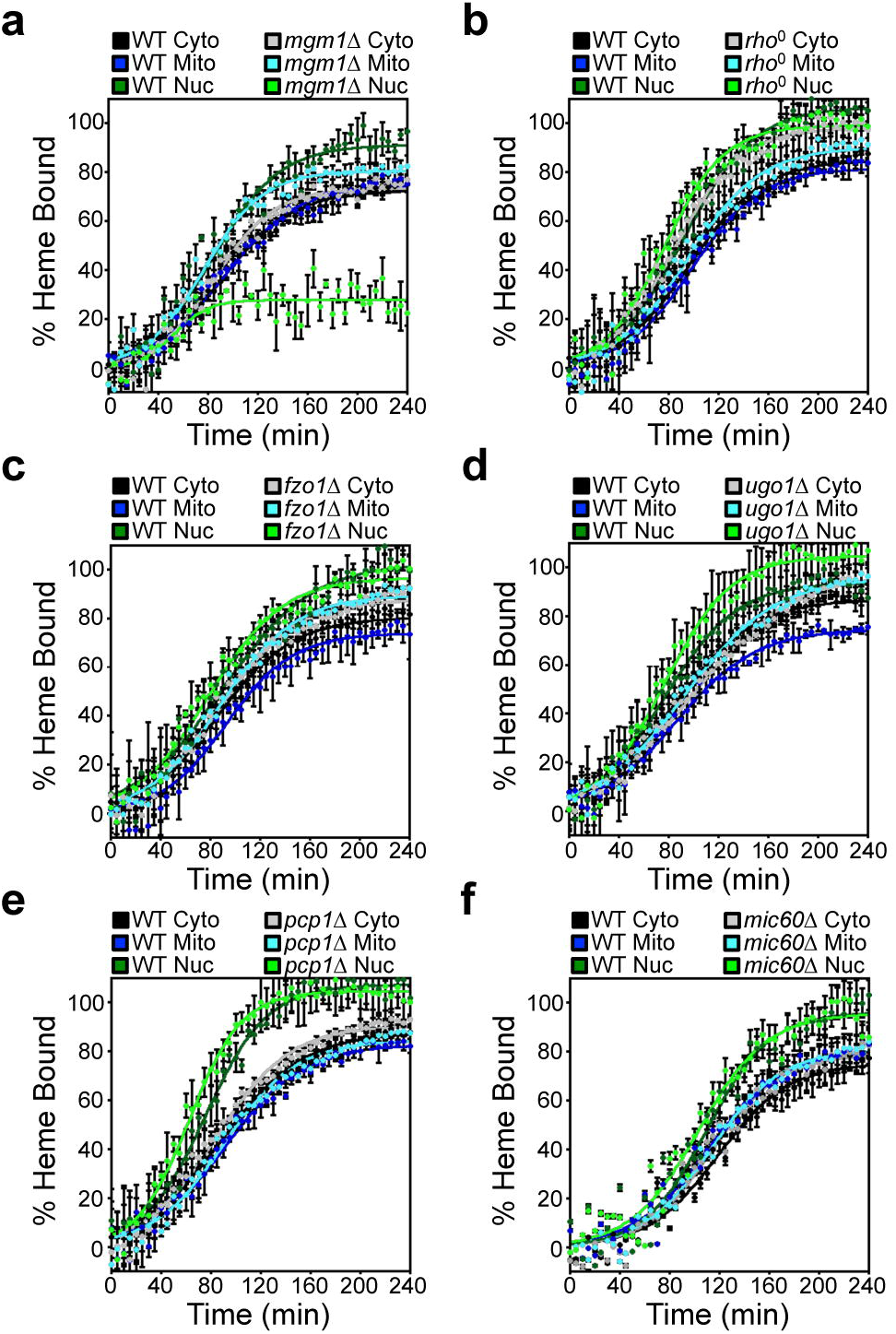
Mgm1 is a positive regulator of mitochondrial-nuclear heme trafficking. Inter-compartmental heme trafficking rates were monitored as in **Fig. 1c** using the SA pulse-chase assay for (**a**) *mgm1*Δ, (**b**) *rho*^0^, (**c**) *fzo1*Δ, (**d**) *ugo1*Δ, (**e**) *pcp1*Δ cells, and (**f**) *mic60*Δ cells. The data represent the mean ± SD of independent triplicate cultures and the kinetic parameters derived from the fits to the data using equation 2 are indicated in **Table S1**.

In order to determine if the nuclear heme trafficking defect observed in *mgm1*Δ cells is specific to Mgm1 or more generally due to ablation of mitochondrial fusion, we tested other essential components of the mitochondrial fusion machinery, including Fzo1 and Ugo1. Interestingly, the *fzo1*Δ (**Fig. 5c**) and *ugo1*Δ (**Fig. 5d**) mutants do not exhibit defects in mitochondrial-nuclear heme trafficking, indicating that general perturbations to mitochondrial fusion do not impact nuclear heme trafficking.

Since the regulation of nuclear heme transport is specific to Mgm1, we tested a mutant with altered Mgm1 function. *pcp1*Δ cells are defective in fusion due to an inability to proteolytically convert l-Mgm1 to its short isoform. As seen in **Fig. 5e**, *pcp1*Δ cells do not exhibit a defect in mitochondrial-nuclear heme trafficking. These data indicate that full-length Mgm1 regulates mitochondrial-nuclear heme transport, but the short Mgm1 isoform is dispensable.

In addition to mediating mitochondrial inner-membrane fusion, Mgm1 is also responsible for maintaining the proper folding and structure of cristae in the inner-membrane (Meeusen et al., 2006). In order to determine if the defect in mitochondrial-nuclear heme trafficking in *mgm1*Δ cells is related to defects in inner-membrane architecture, we determined if heme trafficking was altered in a mutant with a defect in the mitochondrial contact site and cristae organizing system (MICOS). Mic60 is a core component of MICOS and is required for proper inner-membrane folding(Hessenberger et al., 2017). As seen in **Fig. 5f**, *mic60*Δ cells do not exhibit a defect in mitochondrial-nuclear heme trafficking. Altogether, our data strongly suggest that full length Mgm1, but not mitochondrial fusion or inner-membrane architecture *per se*, positively regulates mitochondrial-nuclear heme trafficking (**Fig. 5**).

#### Mitochondrial Fission

In yeast, mitochondrial fission involves recruitment of the GTPase Dnm1 to its receptor Fis1 on the mitochondrial outer-membrane (OM), which is dependent on paralogous adapter proteins Mdv1 and Caf4 (Westermann, 2008). Once assembled and oligomerized around the OM, Dnm1 drives the GTP-dependent constriction and scission of mitochondrial tubules.

Relative to WT cells, the *dnm1*Δ strain exhibits an increase in the rate of heme trafficking to the mitochondrial matrix, cytosol, and nucleus, with the most pronounced effect on nuclear heme trafficking; *k*^NUC^ / *k*^CYTO^ = 1.5 ± 0.1 in *dnm1*Δ cells vs. *k*^NUC^ / *k*^CYTO^ = 1.2 ± 0.1 in WT cells, one-way ANOVA, p < 0.01, n = 3 (**Fig. 6a**). After prolonged growth, 16 hours, heme availability to HS1 in all three compartments is similar between WT and *dnm1*Δ cells (**Fig. S4a**). Additionally, the rate of heme synthesis is unperturbed by loss of Dnm1, indicating that the increase in the rate of heme trafficking is not due to an increase in the rate of heme synthesis (**Fig. S4c**).

**Figure 6.**
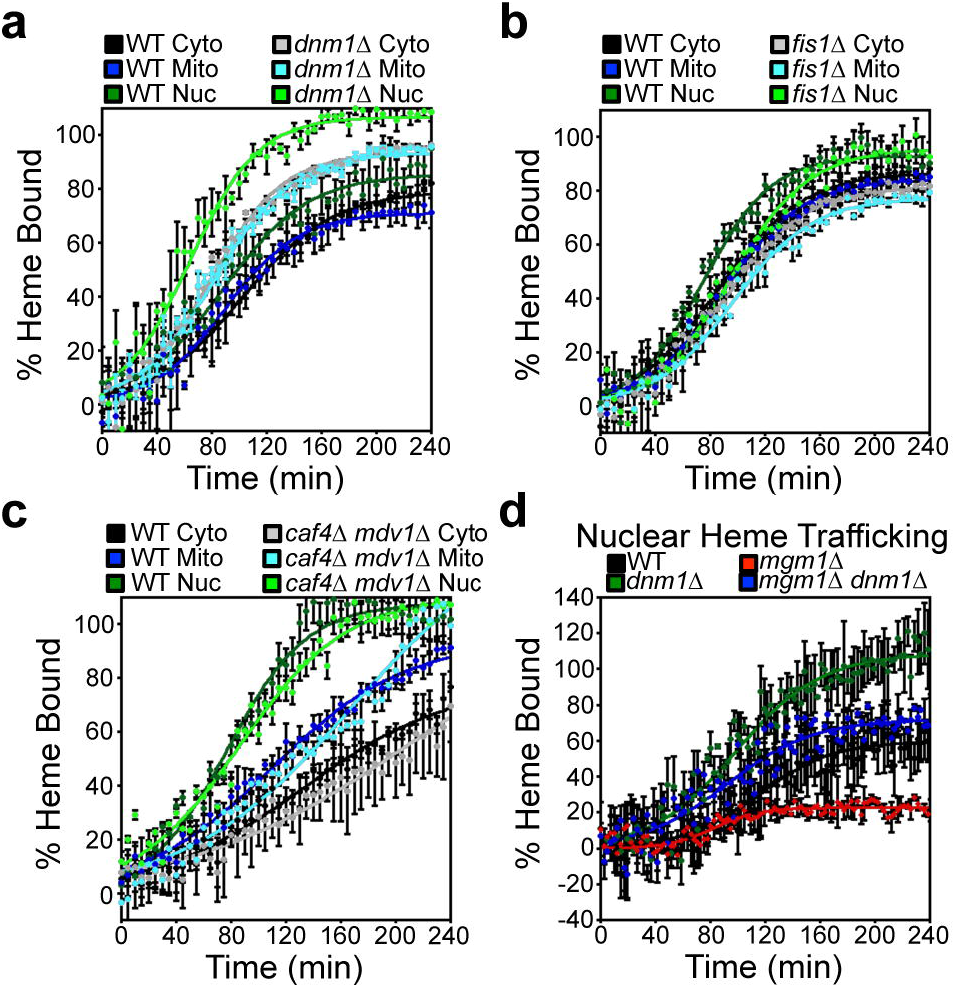
Dnm1 is a negative regulator of mitochondrial-nuclear heme trafficking. Inter-compartmental heme trafficking rates were monitored as in **Fig. 1c** using the SA pulse-chase assay for (**a**) *dnm1*Δ, (**b**) *fis1*Δ, and (**c**) *caf4*Δ *mdv1*Δ cells, and (**d**) *mgm1*Δ *dnm1*Δ cells. Fluorimetry data represent the mean ± SD of three independent cultures and the kinetic parameters derived from the fits to the data using equation 2 are indicated in **Table S1**.

In order to determine if the increased rate of nuclear heme trafficking observed in *dnm1*Δ cells is specific to Dnm1 or more generally due to ablation of mitochondrial fission, we tested other essential components of the mitochondrial fission machinery, including Fis1, Caf4, and Mdv1. Interestingly, *fis1*Δ (**Fig. 6b**) and *caf4*Δ *mdv1*Δ (**Fig. 6c**) mutants do not exhibit perturbations to mitochondrial-nuclear heme trafficking. Altogether, our data indicate that Dnm1, but not mitochondrial fission per se, negatively regulates mitochondrial-nuclear heme trafficking (**Fig. 6**).

We next sought to determine the consequence of ablating both Mgm1 and Dnm1 on mitochondrial-nuclear heme trafficking. Most interestingly, *mgm1*Δ *dnm1*Δ double mutants exhibit WT-like rates of mitochondrial-nuclear heme trafficking (**Fig. 6d**). Therefore, Mgm1 and Dnm1 regulate heme trafficking in opposing directions with similar magnitude.

### Mgm1 and Dnm1 regulate the activation of the nuclear heme-regulated transcription factor Hap1

Given that Mgm1 and Dnm1 are positive and negative regulators of mitochondrial-nuclear heme trafficking, we sought to determine their impact on the activation of the heme-regulated transcription factor Hap1. Heme binding to Hap1 alters its ability to promote or repress transcription of a number of target genes, including *CYC1*, which Hap1 positively regulates (Hanna et al., 2016; Pfeifer et al., 1989; Zhang et al., 1993; Zhang and Hach, 1999; Zhang et al., 1998). In order to probe Hap1 activity, we used a transcriptional reporter that employs the promoter of a Hap1 target gene, p*CYC1*, driving the expression of enhanced green fluorescent protein (eGFP) (Hanna et al., 2016). Cells were first pre-cultured with 500 μM SA for 16 hours and, following this initial growth period, the SA conditioned cells were diluted into fresh media with (+) or without (Δ) 500 μM SA for an additional 4 hours. A parallel set of cells continuously grown without SA (-) was cultured to provide a read out of steady-state Hap1 activity. As demonstrated in **Fig. 7a**, not only is steady-state Hap1 activity greatly diminished in a heme deficient *hem1*Δ mutant and with SA conditioning (+), as expected, loss of Mgm1 results in a nearly 3-fold reduction in Hap1 activity. On the other hand, loss of Dnm1 does not affect basal Hap1 activity (**Fig. 7a**, -). Since basal Hap1 activity in WT cells may already reflect heme saturation of Hap1, it is possible that the effects of the loss of Dnm1 are masked. To address this, we also measured Hap1 activity under non-heme saturating conditions in which Hap1 activity was probed just 4 hours after alleviation of the SA-mediated block in heme synthesis. Under these conditions, *dnm1*Δ cells have increased Hap1 activity and *mgm1*Δ cells exhibit diminished Hap1 activity relative to WT cells (**Fig. 7a**, Δ).

**Figure 7.**
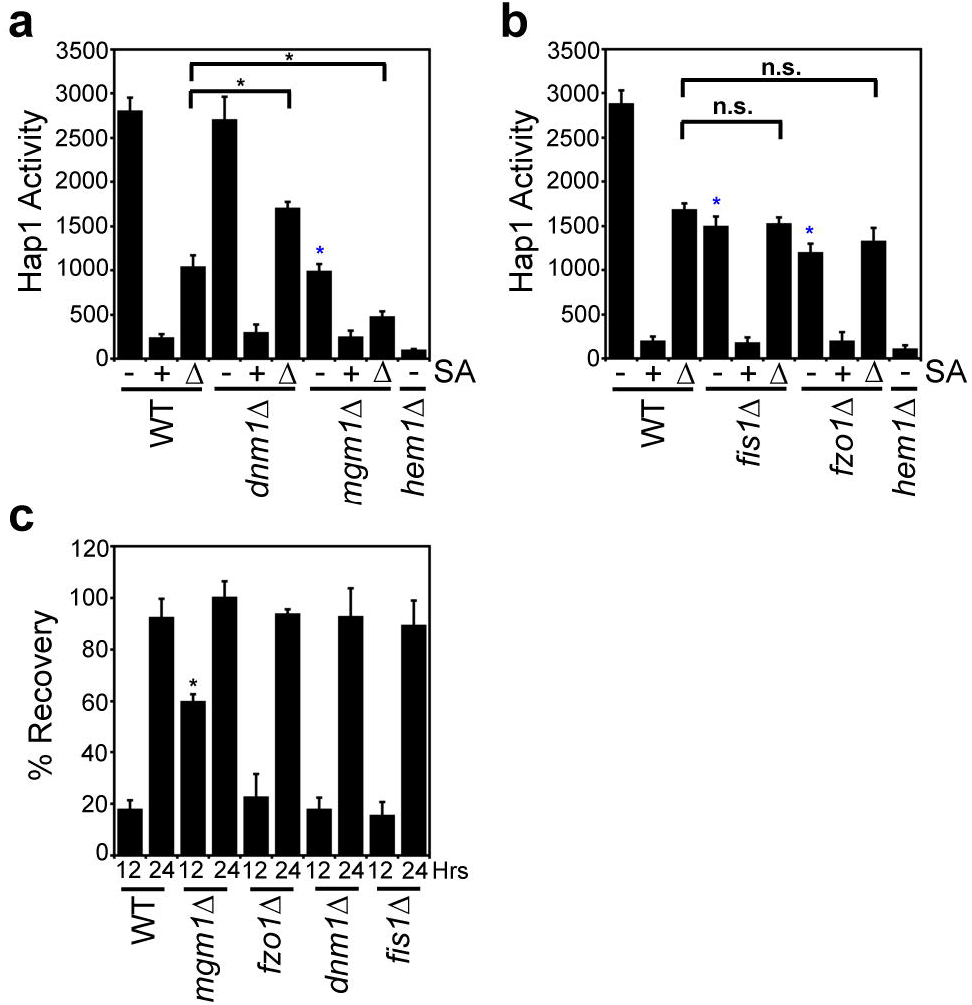
Mitochondrial-nuclear heme trafficking regulates Hap1 activity and sensitivity to new heme synthesis. (**a** and **b**) Hap1p activity in the indicated strains as measured by a transcriptional reporter that uses eGFP driven by the *CYC1* promoter, a Hap1p target gene. We report HAP1 activity in cells that were untreated with succinylacetone (SA) (-), treated with 500 μM SA (+), or were treated with 500 μM SA followed by shifting to media lacking SA for 4 hours (Δ). Fluorimetry data represent the mean ± SD of three biological replicates; *p<0.001 using a one-way ANOVA with Bonferroni’s post-hoc test. Asterisks not associated with an indicated pair-wise comparison were compared to WT untreated samples (-SA). (**c**) Cellular tolerance to new heme synthesis. Cells were pre-cultured overnight with or without 500 μM SA then allowed to recover on media without SA. Sensitivity to new heme synthesis was scored as “% Recovery”, described in Equation 3, by comparing growth to cells continually maintained in media without SA and with SA. Growth data represent the mean ± SD of three biological replicates; *p<0.001 using a one-way ANOVA with Dunnett’s post-hoc test relative to WT cells.

In order to determine if the effects of Hap1 activation by Mgm1 and Dnm1 are unique to these factors, and not general fusion and fission per se, we analyzed Hap1 activity in *fis1*Δ and *fzo1*Δ cells. While steady-state Hap1 activity is diminished in both *fis1*Δ and *fzo1*Δ strains (**Fig. 7b**, -), Hap1 activity probed 4 hours after alleviation of the SA mediated block in heme synthesis results in Hap1 activity that is similar to WT cells (**Fig. 7b**, Δ). Altogether, the positive and negative effects of Mgm1 and Dnm1 on nuclear heme trafficking, respectively, correlate with their effects on heme activation of Hap1, and appear to be distinct from their roles in fusion and fission dynamics (**Fig. 7a** and **7b**, Δ).

We next addressed if Mgm1-and Dnm1-regulated nuclear heme trafficking affected heme-dependent growth of cells. Towards this end, we determined the degree to which heme depleted cells pre-cultured with SA were able to recover in media lacking SA (*G*_ΔSA_) relative to cells continually maintained with (*G*_+SA_) or without SA (*G*_-SA_). As such, the “% Recovery” can be calculated using **Equation 3**:

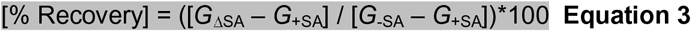

Interestingly, as seen in **Fig. 7c**, after 12 hours of growth, *mgm1*Δ cells are better able to tolerate the re-synthesis of heme compared to WT cells. The superior ability of *mgm1*Δ cells grown with SA to recover in media lacking SA is not phenocopied by another fusion mutant, *fzo1*Δ, suggesting that Mgm1-mediated nuclear heme trafficking may be growth inhibitory. On the other hand, fission mutants, *dnm1*Δ and *fis1*Δ have WT levels of recovery from the alleviation of the SA-mediated block in heme synthesis, presumably because sufficient heme is accessing the nucleus to control growth in a WT-like manner. After 24 hours, all cells achieved maximal growth recovery after re-initiation of heme synthesis (**Fig. 7c**). In total, our data suggests that Mgm1 is not only a positive regulator of mitochondrial-nuclear heme trafficking, but this trafficking is negatively correlated with tolerance to the re-initiation of heme synthesis.

### Heme regulates mitochondrial dynamics

Heme and targets of heme signaling, such as the heme activator proteins, Hap1 and the Hap (Hap2/3/4/5) complex, collectively transcriptionally regulate genes involved in mitochondrial metabolism, including respiration, biogenesis and fission-fusion dynamics (Buschlen et al., 2003; Hon et al., 2005; Knorre et al., 2016; Zhang et al., 2017). We therefore sought to determine if heme itself could regulate mitochondrial fragmentation, a proxy for fission and fusion dynamics. Towards this end, by inhibiting heme synthesis using a 500 μM dose of SA for 4-6 hours, we found that there is a nearly 2-fold increase in mitochondrial fragmentation in SA-treated cells relative to control as early as 4 hours post-treatment (**Fig. 8a** and **8b**). The number of cells with fragmented mitochondria further increases upon prolonged incubation with SA. The 500 μM dose of SA results in a ∼60-80% decrease in intracellular heme (**Fig. 8c**) and alterations in mitochondrial network precede any appreciable changes in mitochondrial membrane potential (**Fig. 8d**). Hence, the SA-dependent increase in mitochondrial fragmentation is consistent with a role for heme in positively influencing mitochondrial fusion and/or negatively regulating mitochondrial fission. Notably, heme-dependent changes in mitochondrial fragmentation alone does not impact heme trafficking dynamics since certain fission and fusion mutants with hyper-fused, *e.g. fis1*Δ cells, or hyper-fragmented, *e.g. fzo1*Δ cells, mitochondria exhibit WT-like rates of heme trafficking.

**Figure 8.**
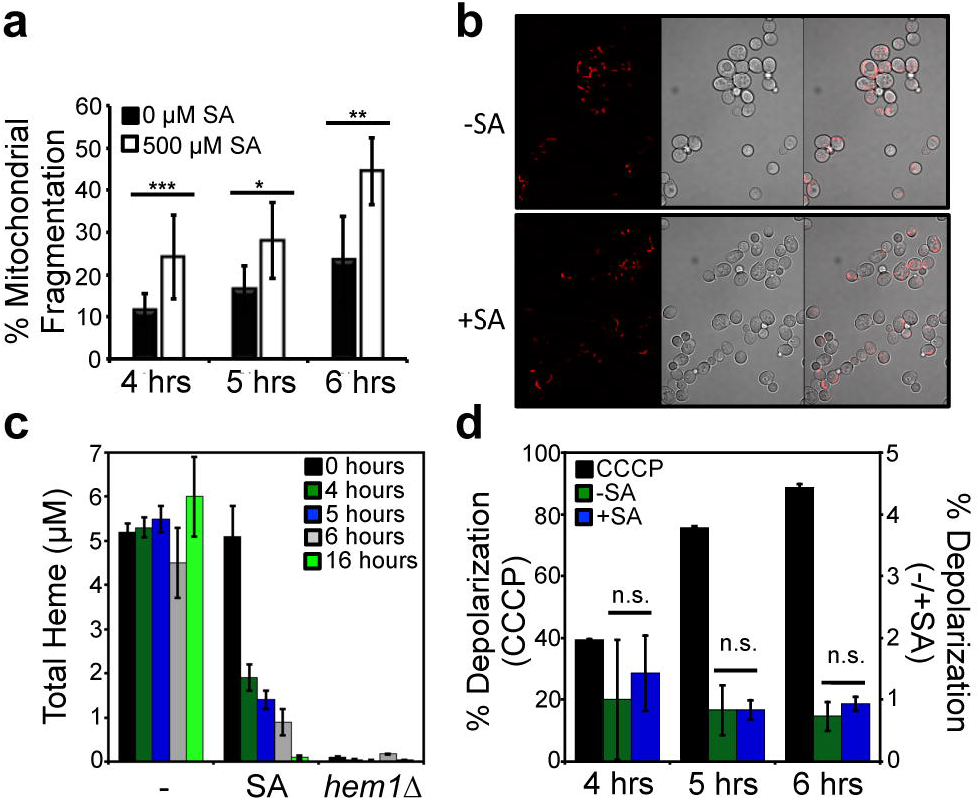
Heme regulates mitochondrial morphology. (**a**) Quantitative analysis of the mitochondrial network in Su9-RFP expressing wild type cells incubated with or without 500 μM succinylacetone (SA) for the indicated periods of time. Mitochondrial fragmentation was quantified as the ratio of the cells displaying punctate or dot-like morphology versus normal tubular mitochondrial networks. Data show mean values ± SD (n=3, with 300-400 cells per biological replicate); *p<0.05, **p<0.01, ***p<0.001 by Student’s *t*-test. (**b**) Representative images of the mitochondrial network from cells used to generate the data in panel **a**. (**c**) Cellular heme levels in wild type cells treated or not with 500 μM SA for an indicated period of time. Heme levels in the *hem1*Δ mutant served as a negative control. Data represent the mean ± SD of three biological replicates. (**d**) Changes in mitochondrial membrane potential in wild type cells treated with 500 μM SA (+SA) or 45 μM uncoupler CCCP (CCCP), or left untreated (-SA) for the indicated periods of time. Cells were stained with 200 nM membrane potential-sensitive dye DiOC_6_ and analyzed by flow cytometry using the FITC channel. Data show the mean values ± SD of 3 independent experiments; n.s., not significant by *t-*test.

## Discussion

Herein, we developed a live-cell assay in yeast to monitor inter-compartmental heme transport dynamics between the mitochondrial IM and the mitochondrial matrix, cytosol, and nucleus in order to identify new aspects of heme trafficking and distribution. *S. cerevisiae* is an unparalleled model system to probe heme distribution dynamics because, unlike many eukaryotes, including mammalian cells, yeast lacking heme can be made viable if supplemented with ergosterol and oleic acid. Thus, heme trafficking and transport dynamics can be monitored without a background associated with pre-existing heme pools. By integrating the heme trafficking dynamics assay with yeast molecular genetics approaches, we have for the first time probed the biodistribution of heme as it is being synthesized in live cells and identified new factors that regulate heme trafficking, including the heme biosynthetic enzyme, ALAS (Hem1) (**Fig. 3**), and GTPases that regulate ERMES, Gem1 (**Fig. 4a**), and mitochondrial fusion, Mgm1 (**Fig. 5a**), and fission, Dnm1 (**Fig. 6**). As discussed below, our results provide fundamental new insights into heme trafficking and signaling.

The heme trafficking kinetics data challenge the current conceptual paradigm for the cellular distribution of mitochondrial heme. The discovery of the first putative mitochondrial heme exporter, Flvcr1b, the only mitochondrial heme transporter identified to date (Chiabrando et al., 2012), has often been taken to imply that heme distribution is sequential, with heme first transported into the cytosol followed by its mobilization to other organelles (Hanna et al., 2017; Reddi and Hamza, 2016; Sweeny et al., 2018). However, we found that heme distribution to the mitochondrial matrix, cytosol, and nucleus occurs virtually simultaneously, suggesting the existence of parallel pathways for heme mobilization to different cellular locales (**Fig. 2** and **3**). Thus, there must be additional factors and mechanisms that distribute heme, in addition to mitochondrial heme transporters that export heme into the cytosol. Our data suggest that GTPases that regulate mitochondrial dynamics and ER contact sites constitute novel components of a new heme distribution network.

We propose that the 25% faster rate of heme trafficking to the nucleus relative to the cytosol or mitochondrial matrix (**Fig. 2f**) is suggestive of heme acting as a mitochondrial-nuclear retrograde signal that enables cells to adapt metabolism and physiology for life with heme and properly functioning mitochondria. Indeed, the activity of the heme regulated nuclear transcription factor, Hap1, correlates with perturbations to mitochondrial-nuclear heme trafficking caused by deletion of Mgm1 or Dnm1, positive and negative regulators of heme trafficking, respectively (**Fig. 7a**). Rather surprisingly, as part of this heme-based retrograde signal, diminished mitochondrial-nuclear heme trafficking in *mgm1*Δ cells correlates with enhanced cellular growth in response to the re-initiation of heme synthesis (**Fig. 7c**). This may be due to the detrimental effects of potentially cytotoxic levels of heme accumulating in the nucleus. Indeed, as an iron containing, redox active, hydrophobic molecule, heme may catalyze spurious redox reactions that promote oxidative damage in the nucleus (Hanna et al., 2017; Reddi and Hamza, 2016). Alternatively, heme may act as an anti-proliferation signal through its ability to act as a transcriptional regulator by binding G-quadruplexes in DNA (Shinomiya et al., 2018; Shumayrikh et al., 2015; Yamamoto et al., 2015) or controlling transcription via Hap1 and the HAP complex (Buschlen et al., 2003; Hon et al., 2005; Zhang et al., 1993; Zhang and Hach, 1999; Zhang et al., 1998). Taken together, our results indicate that the relatively fast rate of nuclear heme acquisition may serve an important role in mitochondrial-nuclear retrograde signaling.

Interestingly, we find that the first enzyme in the heme biosynthetic pathway, ALAS, negatively regulates mitochondrial-nuclear heme trafficking (**Fig. 3**). This result may explain the functional role of a proposed heme biosynthetic supercomplex, or “heme metabolon”, in the IM. The heme metabolon includes the first and last enzymes in the heme synthesis pathway, ALAS and FECH, as well as mitochondrial iron importers and putative heme trafficking factors PGRMC1/2 (Medlock et al., 2015; Piel et al., 2016). The typical biochemical rationale for protein-protein interactions in a biosynthetic pathway usually invokes the requirement for substrate channeling in order to mitigate the dissociation of potentially toxic substrates and ensuring rapid flux. In the case of eukaryotic heme biosynthesis, which is an 8-step process, the first and last two steps occur in the mitochondria with the remaining steps occurring in the cytosol. Thus, the biochemical necessity for an ALAS interaction with FECH is not immediately clear in the context of substrate channeling. Our results suggest that the interaction between ALAS and FECH may be functionally significant from a regulatory perspective because we find that ALAS negatively regulates the flow of heme from FECH in the mitochondrial IM to the nucleus. Since a number of factors are known to regulate ALAS expression and/or mitochondrial import, including glucose, iron, and heme (Keng and Guarente, 1987; Kubota et al., 2016), we propose that the regulation of mitochondrial-nuclear heme trafficking by ALAS may be part of a complex circuit that links nutrient status to heme-based retrograde signaling. However, the precise physiological context and mechanism of ALAS-mediated regulation of mitochondrial-nuclear heme trafficking needs to be further explored.

What is the elusive mechanism of mitochondrial-nuclear heme trafficking? Herein, we identified three conserved GTPases in control of mitochondrial fusion (Mgm1) and fission (Dnm1), and ER-mitochondrial contact sites (Gem1), as regulators of mitochondrial-nuclear heme trafficking. One unifying model consistent with our data is that mitochondrial-ER contact sites facilitate the distribution of heme to the nucleus and other extra-mitochondrial locales. ER tubules facilitate mitochondrial division by constricting mitochondrial tubules so that Dnm1 can oligomerize around the compressed mitochondrial outer-membrane and catalyze the GTP hydrolysis driven scission of mitochondria (Friedman et al., 2011). Gem1 is required to disengage ER-mitochondrial contacts after mitochondrial division(Murley et al., 2013; Nezich and Youle, 2013). Therefore, both *dnm1*Δ and *gem1*Δ cells are expected to - and do - have more stable mitochondrial-ER contact sites (Elbaz-Alon et al., 2014; Murley et al., 2013). Thus, if heme were trafficked through ER-mitochondrial contact sites to reach the ER and organelles contiguous with the ER like the nucleus, one would expect that *dnm1*Δ and *gem1*Δ cells exhibit increased rates of nuclear heme trafficking as was observed (**Fig. 4a** and **6a**).

In accordance with the model just described, we suggest that Mgm1-mediated regulation of mitochondrial-nuclear heme trafficking occurs in a Dnm1 and ERMES-dependent manner. In such a model, Mgm1 promotes mitochondrial-nuclear heme trafficking because of its ability to drive mitochondrial fusion and provide a balancing force to oppose Dnm1-dependent mitochondrial fission. As a consequence, *mgm1*Δ cells would be expected to (and do) exhibit diminished mitochondrial-nuclear heme trafficking since there is no Mgm1 present to counteract Dnm1-mediated fission (**Fig. 5a**). Consistent with this model, we observe a restoration of WT-levels of mitochondrial nuclear heme trafficking in *mgm1*Δ *dnm1*Δ cells (**Fig. 6d**).

However, a number of outstanding issues must be resolved regarding the proposed ERMES-dependent model for mitochondrial-nuclear heme trafficking. First, we do not currently have a means to monitor heme in the ER to more directly assess the impact of ERMES and ER-mitochondrial contact sites on heme trafficking via the ER. Second, it is unclear at this time why other fission and fusion mutants do not exhibit defects in heme trafficking (**Fig 5** and **6**). There may be some specific unknown additional roles of Mgm1 and Dnm1 that allow them to regulate heme trafficking. Third, we do no observe that mutants with defects in essential components of the ERMES complex, including *mdm12*Δ and *mdm34*Δ, exhibit altered mitochondrial-nuclear heme trafficking rates. One explanation is that there are multiple types of ER-mitochondrial contact sites in addition to ERMES that exist. Indeed, a number of factors in a conserved ER membrane protein complex (EMC) have also been found to facilitate physical linkage between the ER and mitochondrial networks (Lahiri et al., 2014). Thus, a heme distribution phenotype may only be observed in multigenic knockout cells defective in both ERMES and ECM factors.

An alternative to the unified model that we present for heme trafficking via ER-mitochondrial contact sites is that each GTPase identified may play mechanistically distinct and unrelated roles in heme trafficking. For instance, the human homolog of Mgm1, OPA1, was previously found to physically associate with FECH in the heme metabolon (Piel et al., 2016). As such, it is possible that Mgm1/OPA1 regulates the heme metabolon and its interactions with putative heme trafficking factors, *e.g.* PGRMC1/2, to effect mitochondrial-nuclear heme trafficking. Dnm1 and Gem1, as lipid-interacting outer-membrane associated GTPases, may play roles in heme sequestration. Binding of lipids such as cardiolipin, can stimulate GTPase activity and Dnm1/Drp1-dependent membrane fission events (Macdonald et al., 2014). Given heme’s lipid-like properties (Reddi and Hamza, 2016), we speculate that heme may be sequestered by Dnm1 or Gem1 in the OM to suppress heme trafficking. Gem1 also has roles in mitochondrial positioning and cytoskeleton anchoring that may contribute to heme distribution dynamics (Koshiba et al., 2011). We are currently exploring the molecular mechanisms underlying Mgm1, Dnm1, and Gem1-mediated heme trafficking and if they cooperate through a common mechanism or act independently of each other.

Our data indicates that factors that impact mitochondrial fission, Dnm1, and fusion, Mgm1, have a profound effect on heme distribution kinetics. Interestingly, our data suggest the reverse may be true as well; namely that heme can impact mitochondrial dynamics (**Fig. 8**). It is interesting to speculate that there may be feedback mechanisms by which heme itself, possibly acting through Mgm1 or Dnm1, can regulate heme distribution dynamics between the mitochondrial network and the nucleus. We are currently probing the underlying mechanisms that govern heme-regulation of mitochondrial fragmentation.

Altogether, the *in vivo* approach to monitor real-time dynamics of inter-compartmental heme trafficking coupled with molecular genetic approaches have uncovered fundamental aspects of the mechanisms underlying heme mobilization and utilization. We expect that the tools and approaches presented herein may be used to probe a number of human diseases, including certain cancers, neurodegenerative disorders, and blood diseases, that are associated with both defects in heme homeostasis and mitochondrial dynamics and membrane contact sites (Atamna and Frey, 2004; Atamna et al., 2002; Wang et al., 2008). Indeed, given that mitochondrial dynamics and contact sites with the endomembrane system are conserved between yeast and man, and the factors that regulate them have homologs or functional analogs between lower and higher eukaryotes (Abrams et al., 2015; Cipolat et al., 2004; MacVicar and Langer, 2016; Schrepfer and Scorrano, 2016; Westermann, 2008), our studies in yeast may be of broad applicability to better understand how membrane and organelle dynamics impacts heme transport and trafficking.

## Materials and Methods

### Yeast Strains, transformations, and growth conditions

*S. cerevisiae* strains used in this study were derived from BY4741 (MATa, *his3*Δ1, *leu2*Δ0, *met15*Δ0, *ura3*Δ0). *fis1*Δ::*Kan*MX4, *dnm1*Δ::*Kan*MX4, *mgm1*Δ::*Kan*MX4, *pcp1*Δ::*Kan*MX4, *ugo1*Δ::*Kan*MX4, *caf4*Δ*::KanMX4*, *mdv1*Δ*::KanMX4* strains were obtained from the yeast gene deletion collection (Thermo Fisher Scientific). We also utilized the previously reported strains, LJ109 (rho^0^) (Reddi and Culotta, 2013) and DH001b-3 (*hem1*Δ::*HIS3*) (Hanna et al., 2016). AR1029-3 (*mdv1*Δ::*Kan*MX4 *caf4*Δ::*HIS3*) was generated by deleting *CAF4* with pAR1047 in *mdv1*Δ::*Kan*MX4 cells. OM232 (*mgm1*Δ::*HIS3*) and OM233 (*dnm1*Δ::*Kan*MX4 *mgm1*Δ::*HIS3*) was generatedby deleting *MGM1* with pAR1051 in WT and *dnm1*Δ*::KanMX4* cells, respectively. All strains were confirmed by PCR, mitochondrial morphology (Fig. S7), and, if derived from the yeast deletion collection, sequencing the unique barcodes flanking the *KanMX4* deletion cassette.

Yeast transformations were performed by the lithium acetate procedure (Gietz and Schiestl, 1991). Strains were maintained at 30° C on either enriched yeast extract (1%) - peptone (2%) based medium supplemented with 2% glucose (YPD), or synthetic complete medium (SC) supplemented with 2% glucose and the appropriate amino acids to maintain selection(Hanna et al., 2016). Cells cultured on solid media plates were done so with YPD or SC media supplemented with 2% agar (Hanna et al., 2016). Selection for yeast strains containing the KanMX4 marker was done with YPD agar plates supplemented with G418 (200 μg/mL) (Hanna et al., 2016). WT cells treated with the heme synthesis inhibitor, succinylacetone (SA), and *hem1*Δ cells were cultured in YPD or SC media supplemented with 50 μg/mL of 5-aminolevulinic acid (ALA) or 15 mg/mL of ergosterol and 0.5% Tween-80 (YPDE or SCE, respectively) (Hanna et al., 2016; Ness et al., 1998). All liquid cultures were maintained at 30 °C and shaken at 220 RPM.

### Plasmids

The *caf4*::*HIS3* disruption plasmid, pAR1047, was generated by first PCR amplifying the upstream (−650 to −129) and downstream (+2110 to +2479) sequences relative to the *CAF4* translational start site, introducing 5’ BamHI / 3’ XhoI and 5’ XbaI / 3’ BamHI restriction sites, respectively. The *CAF4* PCR products were digested with the enzymes indicated and ligated in a trimolecular reaction into the *HIS3* integrating plasmid pRS403 (Sikorski and Hieter, 1989) digested with XhoI and XbaI, resulting in pAR1047. Transformation of yeast strains with pAR1047 linearized with BamHI resulted in deletion of *CAF4* sequences from −128 to +2109.

The *mgm1*::*HIS3* disruption plasmid, pAR1051, was generated by first PCR amplifying the upstream (−532 to −5) and downstream (+2736 to +3023) sequences relative to the *MGM1* translational start site, introducing 5’ BamHI / 3’ XhoI and 5’ XbaI / 3’ BamHI restriction sites, respectively. The *MGM1* PCR products were digested with the enzymes indicated and ligated in a trimolecular reaction into the *HIS3* integrating plasmid pRS403 (Sikorski and Hieter, 1989) digested with XhoI and XbaI, resulting in pAR1051. Transformation of yeast strains with pAR1051 linearized with BamHI resulted in deletion of *MGM1* sequences from −4 to +2735.

Cytosolic, mitochondrial, and nuclear-targeted heme sensors, HS1, were sub-cloned into pRS415 and driven by *ADH*, *TEF*, or *GPD* promoters as previously described (Hanna et al., 2016). The Hap1 reporter plasmid in which eGFP is driven by the *CYC1* promoter was also previously described (Hanna et al., 2016).

### Heme trafficking dynamics assay

*Overview.* Inter-compartmental heme trafficking rates were monitored by: **a.** inhibiting heme synthesis with succinylacetone (SA) in sensor expressing cells; **b.** removing the block in heme synthesis by re-suspending cells into media lacking SA; and **c.** monitoring the time-dependent change in the percentage of heme bound to heme sensor 1 (HS1) localized to the cytosol, nucleus, and mitochondrial matrix upon the re-initiation of heme synthesis. The fractional heme saturation of the sensor can be determined using previously established sensor calibration protocols(Hanna et al., 2016). The % of sensor bound to heme, % Bound, is calculated by determining the sensor eGFP/mKATE2 fluorescence ratio (*R*) under a given test condition relative to the eGFP/mKATE2 fluorescence ratio when the sensor is 100% (*R*_max_) or 0% (*R*_min_) bound to heme as described in **Equation 1** (Hanna et al., 2016).

*R*_min_ is determined by measuring the HS1 eGFP/mKATE2 ratio in parallel cultures that are conditioned with succinylacetone (SA), which inhibits the second enzyme in the heme biosynthetic pathway, ALA dehydratase (ALAD) (Ebert et al., 1979), and *R*_max_ can be determined by permeabilizing cells and adding an excess of heme to saturate the sensor (Hanna et al., 2016). Given HS1 is quantitatively saturated with heme in the cytosol, nucleus, and mitochondria of WT yeast, we typically determine *R*_max_ by measuring the HS1 eGFP/mKATE2 ratio in parallel WT cultures grown without SA (Hanna et al., 2016).

#### Growth for SA-pulse chase assay

HS1-expressing cells were cultured with or without 500 μM SA (Sigma-Aldrich) in SCE-LEU media. Triplicate 5 mL cultures were seeded at an initial optical density of OD_600nm_ = .01-.02 (2-4 x 10^5^ cells/mL) and grown for 14-16 hours at 30 °C and shaking at 220 RPM until cells reached a final density of OD_600nm_ ∼ 1.0 (2 x 10^7^ cells/mL). After culturing, 1 OD or 2 x 10^7^ cells were harvested, washed twice with 1 mL of ultrapure water, and resuspended in 1 mL of fresh SC-LEU media. The cells that were pre-cultured without SA provided HS1 *R*_max_ values. The SA-conditioned cells were split into two 500 μL fractions. One fraction was treated with 500 μM SA to give HS1 *R*_min_ values. The other fraction was not treated with SA so that heme synthesis could be re-initiated to give compartment-specific heme trafficking rates. HS1 fluorescence was monitored on 200 uL of a 1 OD/mL (2 x 10^7^ cells/mL) cell suspension using black Greiner Bio-one flat bottom fluorescence plates and a Synergy Mx multi-modal plate reader. eGFP (ex. 488 nm, em. 510 nm) and mKATE2 (ex. 588 nm, em. 620 nm) fluorescence was recorded every 5 minutes for 4 hours, with the plate being shaken at medium-strength for 30 seconds prior to each read. Background fluorescence of cells not expressing the heme sensors were recorded and subtracted from the eGFP and mKATE2 fluorescence values.

#### Growth for ALA pulse-chase assay

HS1-expressing *hem1*Δ cells were cultured with or without 400 μM 5-aminolevulinic acid (ALA) (Sigma-Aldrich) in SCE-LEU media. Triplicate 5 mL cultures were seeded at an initial optical density of OD_600nm_ = .01-.02 (2-4 x 10^5^ cells/mL) and grown for 14-16 hours at 30 °C and shaking at 220 RPM until cells reached a final density of OD_600nm_ ∼ 1.0 (2 x 10^7^ cells/mL). After culturing, 1 OD or 2 x 10^7^ cells were harvested, washed twice with 1 mL of ultrapure water, and resuspended in 1 mL of fresh SC-LEU media. The cells that were pre-cultured with ALA provided HS1 *R*_max_ values. The cells that were cultured without ALA were split into two 500 μL fractions. One fraction was used to give HS1 *R*_min_ values. The other fraction was treated with 400 μM ALA so that heme synthesis could be initiated to give compartment-specific heme trafficking rates. HS1 fluorescence was monitored on 200 uL of a 1 OD/mL (2 x 10^7^ cells/mL) cell suspension using black Greiner Bio-one flat bottom fluorescence plates and a Synergy Mx multi-modal plate reader. eGFP (ex. 488 nm, em. 510 nm) and mKATE2 (ex. 588 nm, em. 620 nm) fluorescence was recorded every 5 minutes for 4 hours, with the plate being shaken at medium-strength for 30 seconds prior to each read. Background fluorescence of cells not expressing the heme sensors were recorded and subtracted from the eGFP and mKATE2 fluorescence values.

It should be noted that due to evaporation and drying of cell cultures in the plate reader, reliable measurements could not be achieved after 4 hours of continuous monitoring.

### Total heme quantification

Measurements of total heme were accomplished using a fluorimetric assay designed to measure the fluorescence of protoporphyrin IX upon the release of iron from heme as previously described (Michener et al., 2012). For all total heme measurements, ∼1 x 10^8^ cells were harvested, washed in sterile ultrapure water, and resuspended in 500 μL of 20 mM oxalic acid and stored in a closed box at 4 °C overnight (16-18 hours). Next, an equal volume (500 μL) of 2 M oxalic acid was added to the cell suspensions in 20 mM oxalic acid. The samples were split, with half the cell suspension transferred to a heat block set at 95 °C and heated for 30 minutes and the other half of the cell suspension kept at room temperature (∼25 °C) for 30 minutes. All suspensions were centrifuged for 2 minutes on a table-top microfuge at 21000 x g and the porphyrin fluorescence (ex: 400 nm, em: 620 nm) of 200 μL of each sample was recorded on a Synergy Mx multi-modal plate reader using black Greiner Bio-one flat bottom fluorescence plates. Heme concentrations were calculated from a standard curve prepared by diluting 500-1500 μM hemin chloride stock solutions in 0.1 M NaOH into ultrapure water, which was then added back to extra cell samples as prepared above. In order to calculate heme concentrations, the fluorescence of the unboiled sample (taken to be the background level of protoporphyrin IX) is subtracted from the fluorescence of the boiled sample (taken to be the free base porphyrin generated upon the release of heme iron). The cellular concentration of heme is determined by dividing the moles of heme determined in this fluorescence assay and dividing by the number of cells analyzed, giving moles of heme per cell, and then converting to a cellular concentration by dividing by the volume of a yeast cell, taken to be 50 fL (Hanna et al., 2016).

In order to measure heme synthesis rates, exponential phase cells were pre-conditioned with 500 μM SA for 14-16 hours in SCE-LEU media as described above for the heme trafficking dynamics assay. Following this, cells were washed and resuspended in media lacking SA and aliquots of cells were harvested and washed for total heme quantification as described above.

### Hap1 activity

Cells expressing p415-*CYC1*-*eGFP*, or *eGFP* driven by the Hap1 regulated *CYC1* promoter, were cultured in 50 mLs of SCE-LEU in 250 mL Erlenmeyer flasks, with or without 500 μM SA, for 14-16 hours to an optical density of OD_600nm_ ∼ 1.0 (2 x 10^7^ cells/mL). Cells were washed with sterile ultrapure water and diluted into fresh SC-LEU media at an optical density of OD_600nm_ = 0.25 and allowed to grow for an additional 4 hours. The culture that was initially not conditioned with SA remained untreated during the 4-hour growth phase and these cultures represented basal Hap1 activity. The culture that was pre-conditioned with SA was split into two fractions. One fraction was treated with 500 μM SA for the 4-hour growth phase and these cultures served as a negative control, representing minimal Hap1 mediated activation of *CYC1*. The other fraction was not treated with SA for the 4-hour growth phase and these cultures represented Hap1 activity under conditions in which Hap1 was not saturated with heme. After growth for 4 hours, cells were washed in sterile ultrapure water and resuspended in PBS to a concentration of 1 x 10^8^ cells/mL and 100 uL was used to measure eGFP fluorescence (ex. 488 nm, em. 510 nm). Background auto-fluorescence of cells not expressing eGFP was recorded and subtracted from the p415-*CYC1*-*eGFP* expressing strains.

### Isolation of mitochondria and nuclei to confirm heme sensor localization

Mitochondria and nuclei were isolated using yeast mitochondrial (Cat # K259-50) and nuclear (Cat # K289-50) isolation kits from BioVision according to the manufacturer’s instructions. All reagents were supplied by the isolation kits. For isolation of mitochondria, 50 mLs of sensor expressing cells were cultured in SC-LEU in 250 mL Erlenmeyer flasks to a final density of OD_600nm_ = 1.0 (2 x 10^7^ cells/mL). 4 x 10^8^ cells were harvested, washed in ultrapure water, and resuspended in “Buffer A” containing 10 mM DTT for 30 minutes at 30 °C with occasional gentle agitation. Cells were then harvested by centrifugation at 1,500 x g for 5 minutes at room temperature. Cells were then re-suspended in 1 mL of “Buffer B” containing manufacturer provided Zymolyase and incubated at 30 °C with occasional gentle inversion. Spheroplast formation was monitored by diluting 10 uL of the cell suspension into ultrapure water and monitoring the decrease in OD_600nm_ such that it was > 80% less than the initial value, which typically took 30-60 minutes. Spheroplasts were harvested by centrifugation at 1500 x g for 5 minutes, resuspended in “Homogenization Buffer” containing a protease inhibitor cocktail, and lysed with 15 strokes of a Dounce Homogenizer on ice. The lysate was centrifuged at 600 x g for 5 minutes at 4 °C to remove cell debris. The supernatant, which contained the mitochondria, was again centrifuged for 5 minutes at 600 x g to remove additional debris. Finally, mitochondria was harvested by centrifugation at 12000 x g for 10 minutes at 4 °C. The mitochondrial pellet was resuspended in 50 μL of “Storage Buffer”. In order to validate sensor mitochondrial localization, 5% of the total spheroplast fraction, mitochondrial fraction, and the post-mitochondrial fraction (supernatant from the 12000 x g centrifugation step) were electrophoresed on a 14% tris-glycine SDS-PAGE gel and immunoblotted using α-PGK1 (Life Technologies; Cat # PA528612; 1:5000 dilution), a cytosolic marker protein, α-porin (Life Technologies; Cat # 459500; 1:5000 dilution), a mitochondrial marker protein, and α-GFP (Genetex; Cat # GTX30738; 1:5000 dilution) to probe HS1 expression. A goat α-rabbit secondary antibody conjugated to a 680 nm emitting fluorophore (VWR/Biotium; Cat # 89138-520; 1:10000 dilution) was used to probe for PGK1 and GFP. A goat α-mouse secondary antibody conjugated to a 680 nm emitting fluorophore (VWR/Biotium; Cat # 89138-516; 1:10000 dilution) was used to probe for Porin. All gels were imaged on a LiCOR Odyssey Infrared imager (Hanna et al., 2016; Reddi and Culotta, 2013).

For isolation of nuclei, 50 mLs of sensor expressing cells were cultured in SC-LEU in 250 mL Erlenmeyer flasks to a final density of OD_600nm_ = 1.0 (2 x 10^7^ cells/mL). 4 x 10^8^ cells were harvested, washed in ultrapure water, and resuspended in “Buffer A” containing 10 mM DTT for 30 minutes at 30 °C with occasional gentle agitation. Cells were then harvested by centrifugation at 1,500 x g for 5 minutes at room temperature. Cells were then re-suspended in 1 mL of “Buffer B” containing manufacturer provided Zymolyase and incubated at 30 °C with occasional gentle inversion. Spheroplast formation was monitored by diluting 10 uL of the cell suspension into ultrapure water and monitoring the decrease in OD_600nm_ such that it was > 80% less than the initial value, which typically took 30-60 minutes. Spheroplasts were harvested by centrifugation at 1500 x g for 5 minutes, resuspended in 1 mL of “Buffer N” containing a protease inhibitor cocktail, and lysed with 5 strokes of a Dounce Homogenizer on ice. The suspension was incubated for 30 minutes at room temperature with gentle agitation every 3-5 minutes. The lysate was centrifuged at 1500 x g for 5 minutes at 4 °C to remove cell debris. Finally, nuclei were harvested by centrifugation at 20000 x g for 10 minutes at 4 °C. The nuclear pellet was resuspended in 100 μL of “Buffer N”. In order to validate sensor nuclear localization, 5% of the total spheroplast fraction, nuclear fraction, and the post-nuclear fraction (supernatant from the 20000 x g centrifugation step) were electrophoresed on a 14% tris-glycine SDS-PAGE gel and immunoblotted using α-PGK1 (Life Technologies; Cat # PA528612; 1:5000 dilution), a cytosolic marker protein, α-NOP1 (Life Technologies; Cat # MA110025; 1:10000 dilution), a nuclear marker protein, and α-GFP (Genetex; Cat # GTX30738; 1:5000 dilution) to probe HS1 expression. A goat α-rabbit secondary antibody conjugated to a 680 nm emitting fluorophore (VWR/Biotium; Cat # 89138-520; 1:10000 dilution) was used to probe for PGK1 and GFP. A goat α-mouse secondary antibody conjugated to a 680 nm emitting fluorophore (VWR/Biotium; Cat # 89138-516; 1:10000 dilution) was used to probe for NOP1. All gels were imaged on a LiCOR Odyssey Infrared imager (Hanna et al., 2016; Reddi and Culotta, 2013).

### Confirmation of heme sensor localization by microscopy

To confirm mitochondrial or nuclear localization of the sensors, prior to microscopy, 1 x 10^7^ exponential phase cells were incubated with 2.0 μg/mL 4’, 6-diamidino-2-phenylindole (DAPI) (Invitrogen) in SC-LEU media for 45-60 minutes to stain nuclear or mitochondrial DNA (Hanna et al., 2016). Laser scanning confocal microscopy was accomplished on a Zeiss ELYRA LSM 780 Super-resolution Microscope equipped with a 63x, 1.4 numerical aperture oil objective. DAPI was excited at 405 nm and emission was collected using 410-483 nm band pass filters. eGFP was excited with the 488 nm line of an argon ion laser, while mKATE2 was excited using the 594 nm of a HeNe laser line. The 491-588 nm and 599-690 nm band pass filters were used to filter emission for eGFP and mKATE, respectively. Images were collected using Zeiss software and analyzed with ImageJ 1.48v (Rasband, W.S., ImageJ, U. S. National Institutes of Health, Bethesda, Maryland, USA, http://rsb.info.nih.gov/ij/, 1997-2007). Auto-fluorescence of unlabeled cells was subtracted from the DAPI, eGFP, and mKATE2 channels to produce the finalized images in **Figure 1b**.

### Confirmation of mitochondrial morphology in fission and fusion

To visualize the mitochondrial network, 1 x 10^7^ exponential phase cells were incubated with 500 μM Mito Tracker Red CM-H_2_XRos (Thermo Fisher) in SC-LEU media for 45-60 minutes. Confocal microscopy was done on a Zeiss ELYRA LSM 780 Super-resolution Microscope equipped with a 63x, 1.4 numerical aperture oil objective. Mito Tracker Red was excited using the 594 nm of a HeNe laser line and emission was collected using a 415-735 nm band pass filter. Images were collected using Zeiss software and analyzed with ImageJ 1.48v (Rasband, W.S., ImageJ, U. S. National Institutes of Health, Bethesda, Maryland, USA, http://rsb.info.nih.gov/ij/, 1997-2007). The various mitochondrial network morphology types observed are outlined in **Fig. S5a** and ∼50 cells from each strain were analyzed to confirm the expected mitochondrial morphology (**Fig. S5b**). A representative image of the mitochondrial network from each strain is depicted in **Fig. S5c**. Fission and fusion mutants have characteristic elongated or punctate mitochondrial networks, respectively.

### Mitochondrial fragmentation assay

Mitochondrial morphology was assessed in wild-type yeast (W303, *MAT ade2-1 can1-100 his3-11,15 leu2-3,112 trp1-1 ura3-1*) transformed with the pYX142-Su9-RFP plasmid (Frederick et al., 2008). The pYX142-Su9-RFP plasmid expresses a mitochondria-targeted red fluorescent protein (RFP), allowing for visualization of mitochondria using a fluorescence microscope. Cells were grown overnight in synthetic complete (SC) medium supplemented with appropriate nutrients and 2% glucose as the sole carbon source. The cells were then sub-cultured into fresh pre-warmed medium at an OD_600_ of 0.2 for three hours, and then treated with succinylacetone (SA) at a final concentration of 500 μM and cultured for 4, 5, or 6 hours post-SA treatment. Imaging was performed using the Olympus IX81-FV5000 confocal laser-scanning microscope at 543-nm laser line with a 100x oil objective. Images were acquired and processed with Fluoview 500 software (Olympus America). Mitochondrial fragmentation was quantified based on the ratio of the cells displaying punctated or dotted-like morphology versus normal ribbon-like mitochondrial networks. At least 300 cells were analyzed for each sample per replicate.

### Mitochondrial membrane potential assay

The mitochondrial membrane potential was assayed on cells cultured in a similar manner to that described above for the assessment of heme-dependent changes in mitochondrial fragmentation using a BD FACSCanto^TM^II system (Becton Dickinson) as previously described (Pan and Shadel, 2009). After culturing for the indicated times, cells were first washed with phosphate-buffered saline (PBS), and then stained with 200 nM 3,3′-dihexyloxacarbocyanine iodide (DiOC_6_) (Molecular Probes) for 30 min at 30 ^°^C. Finally, the cells were washed twice with PBS and analyzed by flow cytometry using the FL3 channel without compensation. The data were collected, analyzed and plotted using BD FACSDiva software v6.1.1 (Becton Dickinson).

## Acknowledgements

This work was supported by the National Institutes of Health (ES025661 to ARR, GM108975 to OK, DK111653 to AEM), the National Science Foundation (MCB-1552791 to ARR), the Blanchard Professorship (to ARR), Georgia Institute of Technology (to ARR), the U.S. Department of Education GAANN Program (Grant P200A120081 to OMG), and the University of Nebraska Molecular Mechanisms of Disease graduate program (to JVD). We wish to acknowledge the core facilities at the Parker H. Petit Institute for Bioengineering and Bioscience at the Georgia Institute of Technology for use of the shared equipment, services, and expertise.

## References

Abrams, A. J., Hufnagel, R. B., Rebelo, A., Zanna, C., Patel, N., Gonzalez, M. A., Campeanu, I. J., Griffin, L. B., Groenewald, S., Strickland, A. V. et al. (2015). Mutations in SLC25A46, encoding a UGO1-like protein, cause an optic atrophy spectrum disorder. Nat Genet 47, 926–32.

Arosio, P., Knowles, T. P. and Linse, S. (2015). On the lag phase in amyloid fibril formation. Phys Chem Chem Phys 17, 7606–18.

Atamna, H. and Frey, W. H., 2nd. (2004). A role for heme in Alzheimer’s disease: heme binds amyloid beta and has altered metabolism. Proc Natl Acad Sci U S A 101, 11153–8.

Atamna, H., Killilea, D. W., Killilea, A. N. and Ames, B. N. (2002). Heme deficiency may be a factor in the mitochondrial and neuronal decay of aging. Proc Natl Acad Sci U S A 99, 14807–12.

Barr, I., Smith, A. T., Chen, Y., Senturia, R., Burstyn, J. N. and Guo, F. (2012). Ferric, not ferrous, heme activates RNA-binding protein DGCR8 for primary microRNA processing. Proc Natl Acad Sci U S A 109, 1919–24.

Boldogh, I. R., Nowakowski, D. W., Yang, H. C., Chung, H., Karmon, S., Royes, P. and Pon, L. A. (2003). A protein complex containing Mdm10p, Mdm12p, and Mmm1p links mitochondrial membranes and DNA to the cytoskeleton-based segregation machinery. Mol Biol Cell 14, 4618–27.

Burgess, S. M., Delannoy, M. and Jensen, R. E. (1994). MMM1 encodes a mitochondrial outer membrane protein essential for establishing and maintaining the structure of yeast mitochondria. J Cell Biol 126, 1375–91.

Burton, M. J., Kapetanaki, S. M., Chernova, T., Jamieson, A. G., Dorlet, P., Santolini, J., Moody, P. C., Mitcheson, J. S., Davies, N. W., Schmid, R. et al. (2016). A heme-binding domain controls regulation of ATP-dependent potassium channels. Proc Natl Acad Sci U S A 113, 3785–90.

Buschlen, S., Amillet, J. M., Guiard, B., Fournier, A., Marcireau, C. and Bolotin-Fukuhara, M. (2003). The S. Cerevisiae HAP complex, a key regulator of mitochondrial function, coordinates nuclear and mitochondrial gene expression. Comp Funct Genomics 4, 37–46.

Chan, D. C. (2012). Fusion and fission: interlinked processes critical for mitochondrial health. Annu Rev Genet 46, 265–87.

Chen, J. J. (2007). Regulation of protein synthesis by the heme-regulated eIF2alpha kinase: relevance to anemias. Blood 109, 2693–9.

Chiabrando, D., Marro, S., Mercurio, S., Giorgi, C., Petrillo, S., Vinchi, F., Fiorito, V., Fagoonee, S., Camporeale, A., Turco, E. et al. (2012). The mitochondrial heme exporter FLVCR1b mediates erythroid differentiation. J Clin Invest 122, 4569–79.

Cipolat, S., Martins de Brito, O., Dal Zilio, B. and Scorrano, L. (2004). OPA1 requires mitofusin 1 to promote mitochondrial fusion. Proc Natl Acad Sci U S A 101, 15927–32.

DeVay, R. M., Dominguez-Ramirez, L., Lackner, L. L., Hoppins, S., Stahlberg, H. and Nunnari, J. (2009). Coassembly of Mgm1 isoforms requires cardiolipin and mediates mitochondrial inner membrane fusion. J Cell Biol 186, 793–803.

Ebert, P. S., Hess, R. A., Frykholm, B. C. and Tschudy, D. P. (1979). Succinylacetone, a potent inhibitor of heme biosynthesis: effect on cell growth, heme content and delta-aminolevulinic acid dehydratase activity of malignant murine erythroleukemia cells. Biochem Biophys Res Commun 88, 1382–90.

Elbaz-Alon, Y., Rosenfeld-Gur, E., Shinder, V., Futerman, A. H., Geiger, T. and Schuldiner, M. (2014). A dynamic interface between vacuoles and mitochondria in yeast. Dev Cell 30, 95–102.

Frederick, R. L., Okamoto, K. and Shaw, J. M. (2008). Multiple pathways influence mitochondrial inheritance in budding yeast. Genetics 178, 825–37.

Friedman, J. R., Lackner, L. L., West, M., DiBenedetto, J. R., Nunnari, J. and Voeltz, G. K. (2011). ER tubules mark sites of mitochondrial division. Science 334, 358–62.

Gietz, R. D. and Schiestl, R. H. (1991). Applications of high efficiency lithium acetate trasformation of intact yeast cells using single-stranded nucleic acids as carrier. Yeast 7, 253–263.

Hanna, D. A., Harvey, R. M., Martinez-Guzman, O., Yuan, X., Chandrasekharan, B., Raju, G., Outten, F. W., Hamza, I. and Reddi, A. R. (2016). Heme dynamics and trafficking factors revealed by genetically encoded fluorescent heme sensors. Proc Natl Acad Sci U S A 113, 7539–44.

Hanna, D. A., Hu, R., Kim, H., Martinez-Guzman, O., Torres, M. P. and Reddi, A. R. (2018). Heme bioavailability and signaling in response to stress in yeast cells. J Biol Chem 293, 12378–12393.

Hanna, D. A., Martinez-Guzman, O. and Reddi, A. R. (2017). Heme Gazing: Illuminating Eukaryotic Heme Trafficking, Dynamics, and Signaling with Fluorescent Heme Sensors. Biochemistry 56, 1815–1823.

Hessenberger, M., Zerbes, R. M., Rampelt, H., Kunz, S., Xavier, A. H., Purfurst, B., Lilie, H., Pfanner, N., van der Laan, M. and Daumke, O. (2017). Regulated membrane remodeling by Mic60 controls formation of mitochondrial crista junctions. Nat Commun 8, 15258.

Hon, T., Lee, H. C., Hu, Z., Iyer, V. R. and Zhang, L. (2005). The heme activator protein Hap1 represses transcription by a heme-independent mechanism in Saccharomyces cerevisiae. Genetics 169, 1343–52.

Honscher, C., Mari, M., Auffarth, K., Bohnert, M., Griffith, J., Geerts, W., van der Laan, M., Cabrera, M., Reggiori, F. and Ungermann, C. (2014). Cellular metabolism regulates contact sites between vacuoles and mitochondria. Dev Cell 30, 86–94.

Kawano, S., Tamura, Y., Kojima, R., Bala, S., Asai, E., Michel, A. H., Kornmann, B., Riezman, I., Riezman, H., Sakae, Y. et al. (2018). Structure-function insights into direct lipid transfer between membranes by Mmm1-Mdm12 of ERMES. J Cell Biol 217, 959–974.

Keng, T. and Guarente, L. (1987). Constitutive expression of the yeast HEM1 gene is actually a composite of activation and repression. Proc Natl Acad Sci U S A 84, 9113–7.

Knorre, D. A., Sokolov, S. S., Zyrina, A. N. and Severin, F. F. (2016). How do yeast sense mitochondrial dysfunction? Microb Cell 3, 532–539.

Kornmann, B., Currie, E., Collins, S. R., Schuldiner, M., Nunnari, J., Weissman, J. S. and Walter, P. (2009). An ER-mitochondria tethering complex revealed by a synthetic biology screen. Science 325, 477–81.

Kornmann, B., Osman, C. and Walter, P. (2011). The conserved GTPase Gem1 regulates endoplasmic reticulum-mitochondria connections. Proc Natl Acad Sci U S A 108, 14151–6.

Koshiba, T., Holman, H. A., Kubara, K., Yasukawa, K., Kawabata, S., Okamoto, K., MacFarlane, J. and Shaw, J. M. (2011). Structure-function analysis of the yeast mitochondrial Rho GTPase, Gem1p: implications for mitochondrial inheritance. J Biol Chem 286, 354–62.

Kubota, Y., Nomura, K., Katoh, Y., Yamashita, R., Kaneko, K. and Furuyama, K. (2016). Novel Mechanisms for Heme-dependent Degradation of ALAS1 Protein as a Component of Negative Feedback Regulation of Heme Biosynthesis. J Biol Chem 291, 20516–29.

Kumar, S. and Bandyopadhyay, U. (2005). Free heme toxicity and its detoxification systems in human. Toxicol Lett 157, 175–88.

Lackner, L. L. (2019). The Expanding and Unexpected Functions of Mitochondria Contact Sites. Trends Cell Biol 29, 580–590.

Lahiri, S., Chao, J. T., Tavassoli, S., Wong, A. K., Choudhary, V., Young, B. P., Loewen, C. J. and Prinz, W. A. (2014). A conserved endoplasmic reticulum membrane protein complex (EMC) facilitates phospholipid transfer from the ER to mitochondria. PLoS Biol 12, e1001969.

Macdonald, P. J., Stepanyants, N., Mehrotra, N., Mears, J. A., Qi, X., Sesaki, H. and Ramachandran, R. (2014). A dimeric equilibrium intermediate nucleates Drp1 reassembly on mitochondrial membranes for fission. Mol Biol Cell 25, 1905–15.

MacVicar, T. and Langer, T. (2016). OPA1 processing in cell death and disease-the long and short of it. J Cell Sci 129, 2297–306.

Medlock, A. E., Shiferaw, M. T., Marcero, J. R., Vashisht, A. A., Wohlschlegel, J. A., Phillips, J. D. and Dailey, H. A. (2015). Identification of the Mitochondrial Heme Metabolism Complex. PLoS One 10, e0135896.

Meeusen, S., DeVay, R., Block, J., Cassidy-Stone, A., Wayson, S., McCaffery, J. M. and Nunnari, J. (2006). Mitochondrial inner-membrane fusion and crista maintenance requires the dynamin-related GTPase Mgm1. Cell 127, 383–95.

Mense, S. M. and Zhang, L. (2006). Heme: a versatile signaling molecule controlling the activities of diverse regulators ranging from transcription factors to MAP kinases. Cell Res 16, 681–92.

Michener, J. K., Nielsen, J. and Smolke, C. D. (2012). Identification and treatment of heme depletion attributed to overexpression of a lineage of evolved P450 monooxygenases. Proc Natl Acad Sci U S A 109, 19504–9.

Murley, A., Lackner, L. L., Osman, C., West, M., Voeltz, G. K., Walter, P. and Nunnari, J. (2013). ER-associated mitochondrial division links the distribution of mitochondria and mitochondrial DNA in yeast. Elife 2, e00422.

Murley, A. and Nunnari, J. (2016a). The Emerging Network of Mitochondria-Organelle Contacts. Mol Cell 61, 648–53.

Murley, A. and Nunnari, J. (2016b). The Emerging Network of Mitochondria-Organelle Contacts. Mol Cell 61, 648–653.

Ness, F., Achstetter, T., Duport, C., Karst, F., Spagnoli, R. and Degryse, E. (1998). Sterol uptake in Saccharomyces cerevisiae heme auxotrophic mutants is affected by ergosterol and oleate but not by palmitoleate or by sterol esterification. J Bacteriol 180, 1913–9.

Nezich, C. L. and Youle, R. J. (2013). Make or break for mitochondria. Elife 2, e00804.

Nguyen, T. T., Lewandowska, A., Choi, J. Y., Markgraf, D. F., Junker, M., Bilgin, M., Ejsing, C. S., Voelker, D. R., Rapoport, T. A. and Shaw, J. M. (2012). Gem1 and ERMES do not directly affect phosphatidylserine transport from ER to mitochondria or mitochondrial inheritance. Traffic 13, 880–90.

Ogawa, K., Sun, J., Taketani, S., Nakajima, O., Nishitani, C., Sassa, S., Hayashi, N., Yamamoto, M., Shibahara, S., Fujita, H. et al. (2001). Heme mediates derepression of Maf recognition element through direct binding to transcription repressor Bach1. EMBO J 20, 2835–43.

Pan, Y. and Shadel, G. S. (2009). Extension of chronological life span by reduced TOR signaling requires down-regulation of Sch9p and involves increased mitochondrial OXPHOS complex density. Aging (Albany NY) 1, 131–45.

Pfeifer, K., Kim, K. S., Kogan, S. and Guarente, L. (1989). Functional dissection and sequence of yeast HAP1 activator. Cell 56, 291–301.

Piel, R. B., 3rd, Dailey, H. A. Jr. and Medlock, A. E. (2019). The mitochondrial heme metabolon: Insights into the complex(ity) of heme synthesis and distribution. Mol Genet Metab.

Piel, R. B., 3rd, Shiferaw, M. T., Vashisht, A. A., Marcero, J. R., Praissman, J. L., Phillips, J. D., Wohlschlegel, J. A. and Medlock, A. E. (2016). A Novel Role for Progesterone Receptor Membrane Component 1 (PGRMC1): A Partner and Regulator of Ferrochelatase. Biochemistry 55, 5204–17.

Puy, H., Gouya, L. and Deybach, J. C. (2010). Porphyrias. Lancet 375, 924–37.

Raghuram, S., Stayrook, K. R., Huang, P., Rogers, P. M., Nosie, A. K., McClure, D. B., Burris, L. L., Khorasanizadeh, S., Burris, T. P. and Rastinejad, F. (2007). Identification of heme as the ligand for the orphan nuclear receptors REV-ERBalpha and REV-ERBbeta. Nat Struct Mol Biol 14, 1207–13.

Reddi, A. R. and Culotta, V. C. (2013). SOD1 integrates signals from oxygen and glucose to repress respiration. Cell 152, 224–35.

Reddi, A. R. and Hamza, I. (2016). Heme Mobilization in Animals: A Metallolipid’s Journey. Acc Chem Res 49, 1104–10.

Sachar, M., Anderson, K. E. and Ma, X. (2016). Protoporphyrin IX: the Good, the Bad, and the Ugly. J Pharmacol Exp Ther 356, 267–75.

Sassa, S. (2004). Why heme needs to be degraded to iron, biliverdin IXalpha, and carbon monoxide? Antioxid Redox Signal 6, 819–24.

Schipper, H. M., Song, W., Zukor, H., Hascalovici, J. R. and Zeligman, D. (2009). Heme oxygenase-1 and neurodegeneration: expanding frontiers of engagement. J Neurochem 110, 469–85.

Schrepfer, E. and Scorrano, L. (2016). Mitofusins, from Mitochondria to Metabolism. Mol Cell 61, 683–94.

Schuler, M. H., Di Bartolomeo, F., Martensson, C. U., Daum, G. and Becker, T. (2016). Phosphatidylcholine Affects Inner Membrane Protein Translocases of Mitochondria. J Biol Chem 291, 18718–29.

Shadel, G. S. and Seidel-Rogol, B. L. (2007). Diagnostic assays for defects in mtDNA replication and transcription in yeast and humans. Methods Cell Biol 80, 465–79.

Shen, J., Sheng, X., Chang, Z., Wu, Q., Wang, S., Xuan, Z., Li, D., Wu, Y., Shang, Y., Kong, X. et al. (2014). Iron metabolism regulates p53 signaling through direct heme-p53 interaction and modulation of p53 localization, stability, and function. Cell Rep 7, 180–93.

Shinomiya, R., Katahira, Y., Araki, H., Shibata, T., Momotake, A., Yanagisawa, S., Ogura, T., Suzuki, A., Neya, S. and Yamamoto, Y. (2018). Characterization of Catalytic Activities and Heme Coordination Structures of Heme-DNA Complexes Composed of Some Chemically Modified Hemes and an All Parallel-Stranded Tetrameric G-Quadruplex DNA Formed from d(TTAGGG). Biochemistry 57, 5930–5937.

Shumayrikh, N., Huang, Y. C. and Sen, D. (2015). Heme activation by DNA: isoguanine pentaplexes, but not quadruplexes, bind heme and enhance its oxidative activity. Nucleic Acids Res 43, 4191–201.

Sikorski, R. S. and Hieter, P. (1989). A system of shuttle vectors and yeast host strains designed for efficient manipulation of DNA in Saccharomyces ceresiviae. Genetics 122, 19–27.

Sogo, L. F. and Yaffe, M. P. (1994). Regulation of mitochondrial morphology and inheritance by Mdm10p, a protein of the mitochondrial outer membrane. J Cell Biol 126, 1361–73.

Sweeny, E. A., Singh, A. B., Chakravarti, R., Martinez-Guzman, O., Saini, A., Haque, M. M., Garee, G., Dans, P. D., Hannibal, L., Reddi, A. R. et al. (2018). Glyceraldehyde-3-phosphate dehydrogenase is a chaperone that allocates labile heme in cells. J Biol Chem 293, 14557–14568.

Tatsuta, T., Scharwey, M. and Langer, T. (2014). Mitochondrial lipid trafficking. Trends Cell Biol 24, 44–52.

Wang, X., Su, B., Siedlak, S. L., Moreira, P. I., Fujioka, H., Wang, Y., Casadesus, G. and Zhu, X. (2008). Amyloid-beta overproduction causes abnormal mitochondrial dynamics via differential modulation of mitochondrial fission/fusion proteins. Proc Natl Acad Sci U S A 105, 19318–23.

Westermann, B. (2008). Molecular machinery of mitochondrial fusion and fission. J Biol Chem 283, 13501–5.

Williamson, D. H. and Fennell, D. J. (1979). Visualization of yeast mitochondrial DNA with the fluorescent stain “DAPI”. Methods Enzymol 56, 728–33.

Wu, M. L., Ho, Y. C., Lin, C. Y. and Yet, S. F. (2011). Heme oxygenase-1 in inflammation and cardiovascular disease. Am J Cardiovasc Dis 1, 150–8.

Xue, Y., Schmollinger, S., Attar, N., Campos, O. A., Vogelauer, M., Carey, M. F., Merchant, S. S. and Kurdistani, S. K. (2017). Endoplasmic reticulum-mitochondria junction is required for iron homeostasis. J Biol Chem 292, 13197–13204.

Yamamoto, Y., Kinoshita, M., Katahira, Y., Shimizu, H., Di, Y., Shibata, T., Tai, H., Suzuki, A. and Neya, S. (2015). Characterization of Heme-DNA Complexes Composed of Some Chemically Modified Hemes and Parallel G-Quadruplex DNAs. Biochemistry 54, 7168–77.

Yang, Z., Keel, S. B., Shimamura, A., Liu, L., Gerds, A. T., Li, H. Y., Wood, B. L., Scott, B. L. and Abkowitz, J. L. (2016). Delayed globin synthesis leads to excess heme and the macrocytic anemia of Diamond Blackfan anemia and del(5q) myelodysplastic syndrome. Sci Transl Med 8, 338ra67.

Youngman, M. J., Hobbs, A. E., Burgess, S. M., Srinivasan, M. and Jensen, R. E. (2004). Mmm2p, a mitochondrial outer membrane protein required for yeast mitochondrial shape and maintenance of mtDNA nucleoids. J Cell Biol 164, 677–88.

Zhang, L., Bermingham-McDonogh, O., Turcotte, B. and Guarente, L. (1993). Antibody-promoted dimerization bypasses the regulation of DNA binding by the heme domain of the yeast transcriptional activator HAP1. Proc Natl Acad Sci U S A 90, 2851–5.

Zhang, L. and Hach, A. (1999). Molecular mechanism of heme signaling in yeast: the transcriptional activator Hap1 serves as the key mediator. Cell Mol Life Sci 56, 415–26.

Zhang, L., Hach, A. and Wang, C. (1998). Molecular mechanism governing heme signaling in yeast: a higher-order complex mediates heme regulation of the transcriptional activator HAP1. Mol Cell Biol 18, 3819–28.

Zhang, T., Bu, P., Zeng, J. and Vancura, A. (2017). Increased heme synthesis in yeast induces a metabolic switch from fermentation to respiration even under conditions of glucose repression. J Biol Chem 292, 16942–16954.

Zick, M., Duvezin-Caubet, S., Schafer, A., Vogel, F., Neupert, W. and Reichert, A. S. (2009). Distinct roles of the two isoforms of the dynamin-like GTPase Mgm1 in mitochondrial fusion. FEBS Lett 583, 2237–43.

